# Grazers affect the composition of dissolved storage glycans and thereby bacterioplankton composition during a biphasic North Sea spring algae bloom

**DOI:** 10.1101/2022.09.22.509014

**Authors:** Chandni Sidhu, Inga V. Kirstein, Cédric L. Meunier, Johannes Rick, Karen H. Wiltshire, Nicola Steinke, Silvia Vidal-Melgosa, Jan-Hendrik Hehemann, Bruno Huettel, Thomas Schweder, Bernhard M. Fuchs, Rudolf I. Amann, Hanno Teeling

**Author notes:** Corresponding authors: Hanno Teeling, Max Planck Institute for Marine Microbiology, Celsiusstraße 1, 28359 Bremen, Rudolf I. Amann, Max Planck Institute for Marine Microbiology, Celsiusstraße 1, 28359 Bremen, Bernhard M. Fuchs, Max Planck Institute for Marine Microbiology, Celsiusstraße 1, 28359 Bremen. E-mail addresses and telephone numbers of all authors: Chandni Sidhu, Inga V. Kirstein, Cédric L. Meunier, Johannes Rick, Karen H. Wiltshire, Nicola Steinke, Silvia Vidal-Melgosa, Jan-Hendrik Hehemann, Bruno Huettel, Thomas Schweder, Bernhard M. Fuchs, Rudolf I. Amann, Hanno Teeling.

## Abstract

Blooms of marine microalgae play a pivotal role in global carbon cycling. Such blooms entail successive blooms of specialized clades of planktonic bacteria that remineralize algal biomass. We investigated the bacterioplankton response to a bloom in the German Bight in spring 2020. Metagenome sequencing at 30 time-points allowed reconstruction of 251 metagenome-assembled genomes (MAGs), 245 representing as yet uncultured species, while corresponding metatranscriptome sequencing highlighted 50 particularly active MAGs. Together with algae, copepod, protist and bacteria diversity and abundance data in combination with physico-chemical data and antibody-based saccharide measurements, we demonstrate (i) how dissolved primary photoassimilated algal and secondary bacterial storage glycans shape the bacterioplankton community composition, and (ii) how grazing on higher trophic levels determines the release of these abundant glycans. We thus elucidate principles governing how bacterioplankton clades respond to algal blooms and collectively remineralize gigatons of carbon annually on a global scale.

## Introduction

The carbon cycle is the largest biogeochemical cycle. Every year more than 100 Gt of carbon are fixed by photosynthesis in about equal amounts by terrestrial plants and marine algae (Field *et al*., 1998). Marine planktonic uni- to multicellular microalgae (phytoplankton) fix about 45-50 Gt annually (Falkowski *et al*., 1998), of which diatoms alone might fix up to 20 Gt (Mann, 1999). This carbon fixation climaxes during phytoplankton blooms, which can occur in the presence of sufficient solar irradiance and inorganic nutrients. Phytoplankton blooms can reach such massive scales that they are visible by satellite, but are also usually rather short-lived. Bloom termination is often a result of compounding effects, such as self-shading, nutrient exhaustion, viral infections, parasitism by marine fungi, oomycetes and algicidal bacteria, as well as grazing by protists (e.g., flagellates, ciliates) and invertebrate metazoans (e.g., copepods) (Yager *et al*., 2001; Castberg *et al*., 2001; Löder *et al*., 2011).

During blooms, copious amounts of organic material are released to the surrounding seawater, either via algal exudation or cell death, fueling the pools of marine dissolved and particulate organic matter (DOM, POM). A portion of the POM can sink to the deep sea where it can be buried for millennia, while recalcitrant DOM can fuel the oceanic pool of persistent organic matter. However, a substantial portion of the organic matter produced during phytoplankton blooms is rapidly remineralized by marine heterotrophic bacteria in surface waters.

Bloom-associated bacteria have co-evolved with photosynthetic microalgae since the latter emerged during the Precambrian proterozoic period roughly two billion years ago (Sergeev *et al*., 2002), including the more recent diatoms that emerged during the Permian-Triassic extinction event about 250 million years ago (Benoiston *et al*., 2017). These bacteria constitute tight-knit communities that collectively decompose algal biomass. Many partaking species are outright specialists that target only specific classes of algal biomass, a strategy which minimizes competition and concomitantly maximizes efficiency (resource partitioning).

Polysaccharides (glycans) represent a major class of algal biomass, used for instance as intracellular energy storage or found in cell matrices and cell walls. Total contents depend on microalgal species and physiological state, and can reach up to 90% of the dry weight (Myklestad, 1974). In addition, algae can exudate glycan-rich transparent exopolymer particles (TEP), especially when under stress. Many algal polysaccharides are anionic, often by sulfation, and have no counterparts in terrestrial plants. Simple algal polysaccharides consist of a sole monosaccharide and few linkage types, such as for example laminarin, a helical polysaccharide composed of glucose monomers with a β-1,3-linked backbone and occasional β-1,6-linked branches. Laminarin serves as storage of photoassimilated glucose in diatoms and therefore is one of the most abundant polysaccharides on Earth (Becker *et al*., 2020). Other algal polysaccharides are structurally more complex and involve numerous monosaccharides, linkage types and secondary modifications. Knowledge on respective structures is sparse, in particular for planktonic microalgae.

Polysaccharide decomposition requires various specifically adapted carbohydrate-active enzymes (CAZymes) belonging to different glycoside hydrolase (GH), polysaccharide lyase (PL), and carbohydrate esterase (CE) families (Cantarel *et al*., 2009). In polysaccharide-degrading bacteria, genes for decomposition and uptake of dedicated polysaccharides are usually organized in operon-like polysaccharide utilization loci (PULs) that allow inferences about the chemical nature of the target polysaccharide substrate. PUL genes can account for a considerable proportion of genes in specialized bacterial clades (e.g. Kappelmann *et al*., 2019). This constitutes a considerable genetic investment, which entails that resource partitioning is particularly pronounced among polysaccharide-degrading bacteria.

We have analyzed the response of planktonic bacteria (bacterioplankton) to spring phytoplankton blooms at Helgoland Roads (German southern North Sea) since 2009 with increasing detail (Teeling *et al*., 2012; Teeling *et al*., 2016; Francis *et al*., 2019; Krüger *et al*., 2019). Suitable conditions for these mostly diatom-dominated blooms usually occur from mid-March to the beginning of April. Initially inorganic nutrients are usually abundant and predator abundances are low, which allows algae and bacteria to grow with few restrictions. This largely bottom-up controlled phase is often characterized by swift successions of distinct algae and bacterioplankton taxa (Teeling *et al*., 2016). The latter usually represent about >99% of the total bacteria in the water column (Heins *et al*., 2021). As blooms progress, parasites and predators start to catch up. Top-down pressure from infections and grazing increases and, together with nutrient depletion, ultimately these blooms are terminated after a few weeks.

Phytoplankton blooms are highly dynamic events. Substantial changes in both algae and bacterioplankton community composition can happen within a day or two. Frequent sampling is thus required in order to disentangle what drives bacterioplankton dynamics. Highly-resolved bacterioplankton community composition analysis is also required, since some clades comprise species with considerable functional diversity, whereby the genus *Polaribacter* is a prime example (Avcı *et al*.,2020). This diversity is usually not well addressed with partial 16S rRNA gene amplicon sequencing in combination with short-read-based metagenomics - methods that were common until recently. Finally, high time-resolution, deep expression analysis is a precondition to capture key metabolic processes of distinct clades over a bloom’s progression.

Here we present a study of the 2020 spring phytoplankton bloom at Helgoland Roads. Over a 90-day period we sampled and sequenced bacterioplankton metagenomes and -transcriptomes at 30 dates. With PacBio Sequel II long read sequencing, we obtained 251 representative metagenome-assembled genomes (MAGs). Subsequent mapping of deeply sequenced Illumina-based short-read metatranscriptomes allowed the identification of active community members and their metabolisms with high resolution and thereby the designation of distinct MAGs to individual bloom phases. Together with *in situ* saccharide measurements, we demonstrate that diatom laminarin and bacterial α-glucan storage polysaccharides were the most prominent dissolved polysaccharides. We show that peaks in the concentrations of both of these glycans induced blooms of specialized bacterioplankton clades with distinct polysaccharide niches. Furthermore, our data suggest that changes in the availabilities of both storage polysaccharides were caused by interdependent changes in protist and copepod grazing patterns.

## Results

### The 2020 phytoplankton bloom was biphasic

The 2020 spring bloom started around March 24^th^ and lasted until the end of May. It consisted of two distinct bloom phases with an inflection point around April 23^rd^ (**Fig. 1**). The first bloom phase started with a sevenfold increase in chlorophyll *a* from ~1 to ~7 μg per liter within just two days and went through a total of three maxima. The combined algal biovolume during this phase was dominated by the large centric diatom species *Ditylum brightwellii* (**Fig. 1A**), with minor contributions from *Phaeocystis* sp. (*Haptophyta*, class *Prymnesiophyceae*), unspecified *Dinophyceae*, as well as *Cerataulina pelagica, Chaetoceros* sp.*, Guinardia delicatula*, and *Thalassiosira rotula* centric diatoms (**Fig. 1B**) *. D. brightwellii* numbers declined towards the end of the first phase with chlorophyll *a* declining to ~2 μg per liter, passing into the second phase. This phase was characterized by rapid proliferation of *Chaetoceros* sp. and in particular large *C. pelagica* centric diatoms, which resulted in an initial chlorophyll *a* increase from below 2 to above 9 μg per liter within five days. During the second phase, also *Phaeocystis* sp. and *Dinophyceae* numbers started to increase, but did not reach notable proportions of the total algal biovolume (**Table S1)**. At the same time, the inorganic nutrients silicate, phosphate, ammonium, and nitrate became scarce (**Fig. S1C,D**, **Table S1**) and the bloom went into decline from about mid-May on. During this terminal phase, *Dinophyceae* numbers decreased less rapidly than diatom numbers, probably because the former do not depend on the silicate which the latter need for frustule formation. At the end of May the bloom was over and algal cell numbers were almost down to pre-bloom levels.

**Fig. 1.**
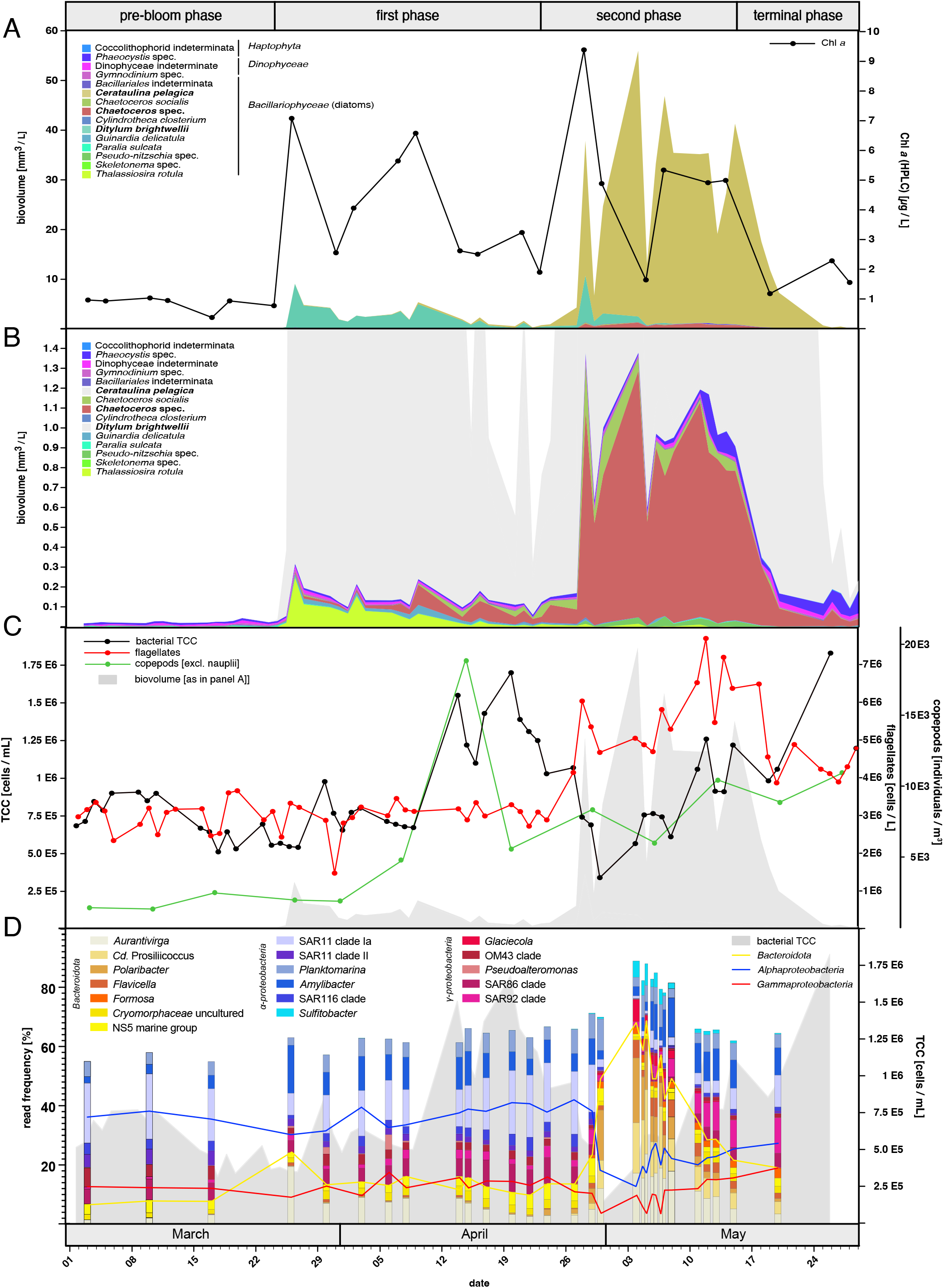
Major phases of the spring phytoplankton bloom at Helgoland Roads during March to May 2020: **A.** Biovolumes of abundant phytoplankton taxa and chlorophyll *a* concentration as measured by HPLC (black line). **B.** Biovolumes of relatively less abundant algal taxa and chlorophyll a concentration. **C.** DAPI-based bacterial total cell counts (TCC; black line), flagellates counts (red line) and counts of copepods (excluding *Copepoda nauplli;* green line). Estimated phytoplankton biovolumes are plotted as a grey area in the background. **D.** Abundances of the 18 most abundant bacterial clades as assessed by metagenomic 16S rRNA gene frequencies. The sum of bacteroidetal, alpha- and gammaproteobacterial clades among these top 18 clades are depicted by lines. Bacterial total cell counts are represented as grey area in the background.

During the first bloom phase, flagellate numbers remained low at an almost constant 2-3*10^6^ per L, indicating substantial grazing on flagellates from higher trophic levels (**Fig. 1C**). Microscopic observations confirmed a pronounced copepod bloom during the second half of this phase that was dominated by *Paracalanus* species (**Table S2**). The copepod bloom started around April 1^st^ and peaked around April 15^th^ (**Fig. 1C**). Copepod grazing most likely caused the observable decrease in algae (**Fig. 1A**), and was accompanied by a pronounced increase in bacterioplankton with total cell counts (TCC) from around 0.5 to 1.7*10^6^ per mL (**Fig. 1C**). It is known that copepod sloppy feeding and excretion increase substrates available to bacteria (e.g., Vargas *et al*., 2007). The decrease in algae abundances likely caused copepod numbers to subside after April 15^th^. The resulting diminished grazing pressure and increased bacterial abundances probably allowed flagellate numbers to double. This, together with a lowered substrate availability due to the decreased algae abundances, likely caused an about four-fold drop in bacterial cell counts to <0.4*10^6^ per mL at the bloom’s inflection point (**Fig. 1C**). During the second phytoplankton bloom phase, which was characterized by much higher overall algae numbers and thus substrate availability, bacterioplankton numbers recovered and increased and reached about 1.8*10^6^ per mL on May 26^th^. Likewise, flagellate abundances increased gradually throughout the second bloom phase, while copepod numbers increased only little (**Fig. 1C**).

On April 1^st^/2^nd^ and at the onset of the second phase on April 22^nd^/23^rd^ inorganic ammonium and phosphate concentrations spiked almost threefold to about 5.1-6.8 μM and 1.6-1.8 μM, respectively. Both spikes were accompanied by 2-3% drops in salinities (**Fig. S1A,D**), indicating incursions of nutrient-rich coastal waters, likely from the coastal Weser and Elbe river estuaries (~50 km distance). On 29^th^/30^th^ April strong rainfall (3.3 to 5.3 L/m^2^) coincided with another decrease in salinity (**Table S1**) and a first peak of *C. pelagica* and *Chaetoceros* species. Besides an initial top-down control, the second weather event at the bloom’s inflection point might have shaped the biphasic nature of the 2020 Helgoland spring bloom.

### 16S rRNA gene frequencies revealed that few flavobacterial clades dominated the second bloom phase

We sampled 30 time points throughout the course of the bloom and sequenced corresponding metagenomes on a PacBio Sequel II (long-read HiFi mode). Unassembled reads contained 325,737 near full-length 16S rRNA gene sequences (average: 1,269 bp) that were used for diversity analyses (**Fig. 1D**, **Table S3 and S4**). Among *Alphaproteobacteria*, SAR11 Ia clade sequences dominated during the pre-bloom and first-bloom phases, reaching up to 22.1% on March 10^th^ and 21.4% on April 20^th^. Likewise, SAR11 clade II sequences were confined to the pre-bloom and first bloom phases and peaked with 5.1% also on March 10^th^. In contrast, *Amylibacter* sequences were detected during all bloom phases reaching up to 16.2% on March 26^th^ and 13.3% on May 7^th^ (9 PM sample). A similar pattern was found for *Planktomarina* with a maximum of ~11% on April 30^th^. *Sulfitobacter* sequences were associated with the second and terminal bloom phases and reached up to ~8%, while SAR116 sequences never exceeded ~5% at any time point.

Sequences of abundant *Bacteroidota* affiliated with well-known bloom-associated *Flavobacteriaceae* clades, such as *Aurantivirga* (Krüger *et al*., 2019), *Cd*.Prosiliicoccus (Francis *et al*., 2019) and *Polaribacter* (Avcı *et al*., 2020). Flavobacterial clades ramped up at the bloom’s inflection point and dominated in the second bloom phase, topping out at 56% on May 4^th^. *Flavobacteriia* relative numbers declined after May 8^th^ during the bloom’s terminal phase. The most abundant flavobacterial clade *Aurantivirga* peaked with 19% of the bacterial relative abundance on March 26^th^ during the first phase, and with ~23% on May 5^th^ during the second phase. In contrast, *Polaribacter* were only detected in the second bloom phase, peaking with 21.7% relative abundances on May 5^th^. *Flavicella* members were present during the second and terminal-bloom phases and reached up to 6.5% on May 6^th^. The NS4, NS5, NS2b, NS3a, and NS9 clades (Alonso *et al*., 2007) occurred during the pre- and first-bloom phases. Sequences of other *Bacteroido*ta clades such as *Formosa*, *Fluviicola, Marinoscillum, Aquibacter*, and uncultured *Cryomorphaceae* were found during all bloom phases with proportions below 5%.

The most prominent *Gammaproteobacteria* were members of the SAR92 clade, in particular during the terminal bloom phase with up to ~12% relative abundance on May 20^th^. In contrast, SAR86 sequences were present mostly during the pre- and second bloom phases, and during the terminal phase after May 8^th^. Members of the OM182 clade followed a similar pattern, but relative abundances never exceeded ~3%. *Methylophilaceae* (OM43 clade, reclassified from *Beta-* to *Gammaproteobacteria* in GTDB (Parks *et al*., 2022)) were also restricted to the pre- and first bloom phases. *Glaciecola* sequences were constricted to the second bloom phase with a maximum of ~8% on May 4^th^. *Pseudoalteromonas* 16S rRNA gene sequences were found in the first bloom phase with a maximum of ~5% on April 6^th^. Sequences of *Luminiphilus* OM60 (NOR5 clade) were found in all bloom phases, but with low frequencies. In contrast, *Cd*. Thioglobus (SUP05 clade) sequences were confined to the pre-bloom phase before March 26^th^.

Sequences of actinobacteriotal *Cd*. Actinomarina were restricted to the pre- and first bloom phases with a maximum of ~5% relative abundance on March 10^th^. Marine group II *Archaea* sequences were low in abundance throughout the bloom, with highest relative abundances of ~11% during the pre-bloom phase on March 26^th^.

### Abundant clades were confirmed by CARD-FISH analyses

Microscopic cell counting with fluorescently labeled CARD-FISH (catalyzed reporter deposition-fluorescence *in situ* hybridization) probes (**Table S5**) confirmed *Aurantivirga* (probe AUR452) and *Polaribacter* (probe POL740) as the dominant clades within the *Bacteroidota* with maximum relative abundances of 18.3% (112,000 cells/mL) and 17.7% (100,000 cells/mL) on May 8^th^ and 4^th^, respectively (**Fig. 2A**). Overall abundances of both clades were highest during the second bloom phase, but seldom exceeded 5% during other phases. This was the case when *Aurantivirga* peaked to slightly above 5% relative abundance on March 26^th^/27^th^ (~27,000 cells/mL). This peak coincided with the initial *D. brightwellii* peak, suggesting a rapid response of *Aurantivirga* to the proliferation of this diatom*. Cd*. Abditibacter (Grieb *et al*., 2020; probe Vis6-814), on the other hand, had low overall abundances, ranging from 0.1% to 3.4% throughout the bloom. Towards the end of the second phase into the bloom’s terminal decline, relative abundances of dominating *Bacteroidota* declined, whereas those of the gammaproteobacterial SAR92 clade (probe SAR92-627) gradually increased to a maximum of 10.7% (105,000 cells/mL) on May 19^th^ (**Fig. 2B**). Members of the gammaproteobacterial SAR86 clade (probe SAR86-1245) exhibited higher relative abundances during the first than the second bloom phase, but values never exceeded 8%. The gammaproteobacterial OM182 clade (probe OM182-707) showed an almost uniform distribution throughout the bloom, with values mostly ranging below 4.5%. Both, SAR86 and OM182 relative abundances tanked below 1% on May 4^th^ and recovered afterwards. Such a minimum was not observed for the SAR92.

**Fig. 2.**
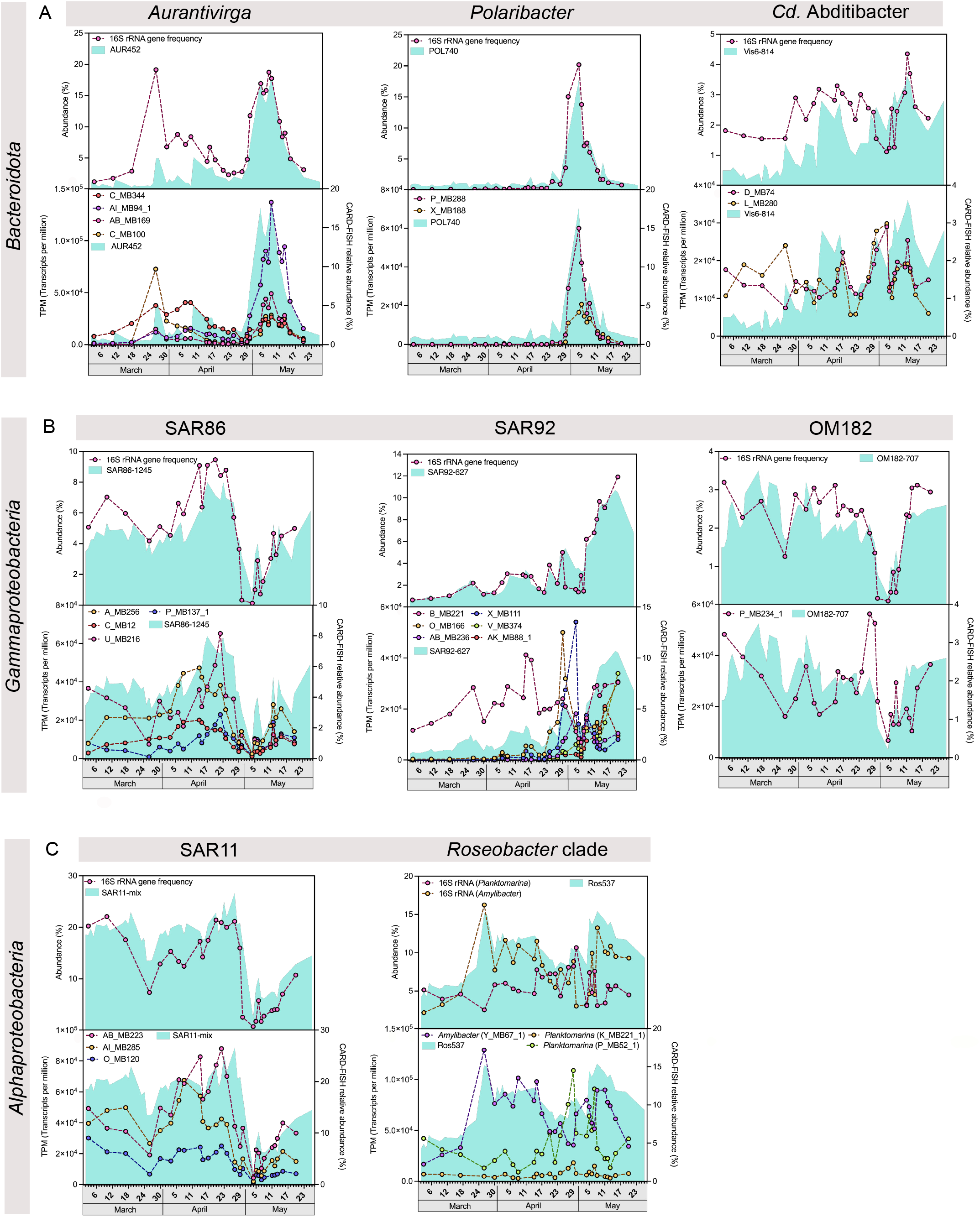
Upper panels: Abundance of various abundant clades as assessed by CARD-FISH (turquoise areas) and by metagenomic 16S rRNAs gene frequencies (lines). Lower panels: Abundances as assessed by CARD-FISH (turquoise areas) and expression patterns of corresponding metagenome assembled genomes as assessed by metatranscriptomics (lines) expressed as transcripts per million (TPM) **A.** *Bacteroidota*: *Aurantivirga* (probe AUR452), *Polaribacter* (probe POL740) and *Cd*. Abditibacter (probe Vis6-814); **B.** *Gammaproteobacteria*: SAR86 clade (probe SAR86-1245), SAR92 clade (probe SAR92-627) and OM182 clade (probe OM182-707); **C.** *Alphaproteobacteria:* SAR11 clade (probe SAR11-mix) and RCA clade *Planktomarina* and *Amylibacter* (probe Ros537).

Corroborating metagenome 16S rRNA gene frequencies, SAR11 (probe SAR11-mix) dominated during the pre-bloom and both major bloom phases with relative abundances ranging from 18.6% (127,000 cells/mL) on March 2^nd^ to 26.6% (286,000 cells/mL) on April 27^th^ (**Fig. 2C**). Like the gammaproteobacterial SAR86 and OM182 clades, SAR11 also suddenly dropped to 2.1%relative abundance on May 4^th^ with a slow recovery afterwards that lasted throughout the bloom’s terminal phase. Members of the abundant *Roseobacter* clade (probe Ros537) were more uniformly distributed, with peaks in relative abundance of around ~15% (~10,000 cells/mL) on March 26^th^/27^th^ (initial *D. brightwellii* peak) during the first bloom phase, and on May 8^th^ during the second bloom phase. Abundant members comprised *Amylibacter* and *Planktomarina*, with *Amylibacter* being abundant throughout the bloom, and *Planktomarina* thriving during the second bloom phase (**Fig. 2C**).

### High-quality representative MAGs could be obtained for most abundant clades

Automatic binning of all 30 individual metagenomes yielded 10,950 initial bins, and 11,071 bins after manual refinement (**Table S6, Fig. S2A**). Of these 1,648 had >70% completeness and <5% contamination estimates (**Fig. S2B**). Dereplication at 0.95 ANI reduced this set to 251 representative, non-redundant MAGs, 165 of which had >90% completeness and <5% contamination estimates. According to MIMAG standards (Bowers *et al*., 2017), 158 MAGs were classified as high-quality MAGs. A total of 140 MAGs consisted of ≤10 contigs, and 76 even had a no predicted contamination, substantiating the dataset’s high quality (**Fig. S2C**).

GTDB taxonomic affiliation revealed that 97.2% of the MAGs belonged to *Bacteria* and 2.8% to *Archaea*. On phylum level, *Proteobacteria* (48.6%) dominated, followed by *Bacteroidota* (36.3%). The remaining 12.3% belonged to *Actinobacteriota, Verrucomicrobiota, Planctomycetota, Marinisomatota, Myxococcota* and *Campylobacterota* (**Table S7**). These representative MAGs co-related well with the overall abundant clades detected by CARD-FISH as well as 16S rRNA gene frequencies.

### Few particularly active MAGs dominated overall expression

Mapping of reads from all 30 transcriptomes onto the complete metagenome dataset recruited 71.5% of the transcripts, and onto the 251 representative MAGs 41.4% of the transcripts. During the pre-bloom, gammaproteobacterial and alphaproteobacterial MAGs dominated with 34.5% (349,824) and 32.1% (320,829) of the total transcripts per million (TTPM), respectively **(Fig. S3**). At the onset of the first bloom phase, relative activities of *Bacteroidia* and *Poseidonia* A (*Thermoplasmatota*) MAGs increased notably and peaked on March 26^th^ during the first *D. brightwellii* peak with 36.3% (363,562) and 20.4% (203,861) of the TTPM, respectively. During the second-bloom phase after April 27^th^, relative activities of alphaproteobacterial MAGs decreased and reached a minimum of 18.0% (179,681) of the TTPM on May 13^th^. Simultaneously, relative activities of *Bacteroidia* MAGs increased during the second-bloom phase and even dominated a larger part of this phase with a maximum of 53.2% (532,164) of the TTPM on May 8^th^ (**Fig. S3**, **Table S8**). The terminal phase lastly was characterized by a decrease in *Bacteroidota* and an increase in *Gammaproteobacteria* transcripts.

As the bulk of metatranscriptome reads mapped to a small number of particularly active MAGs, we confined our analysis to the 50 topmost expressed MAGs representing 30.3% of all transcripts (**Figs. 3** and **S4**).

**Fig. 3.**
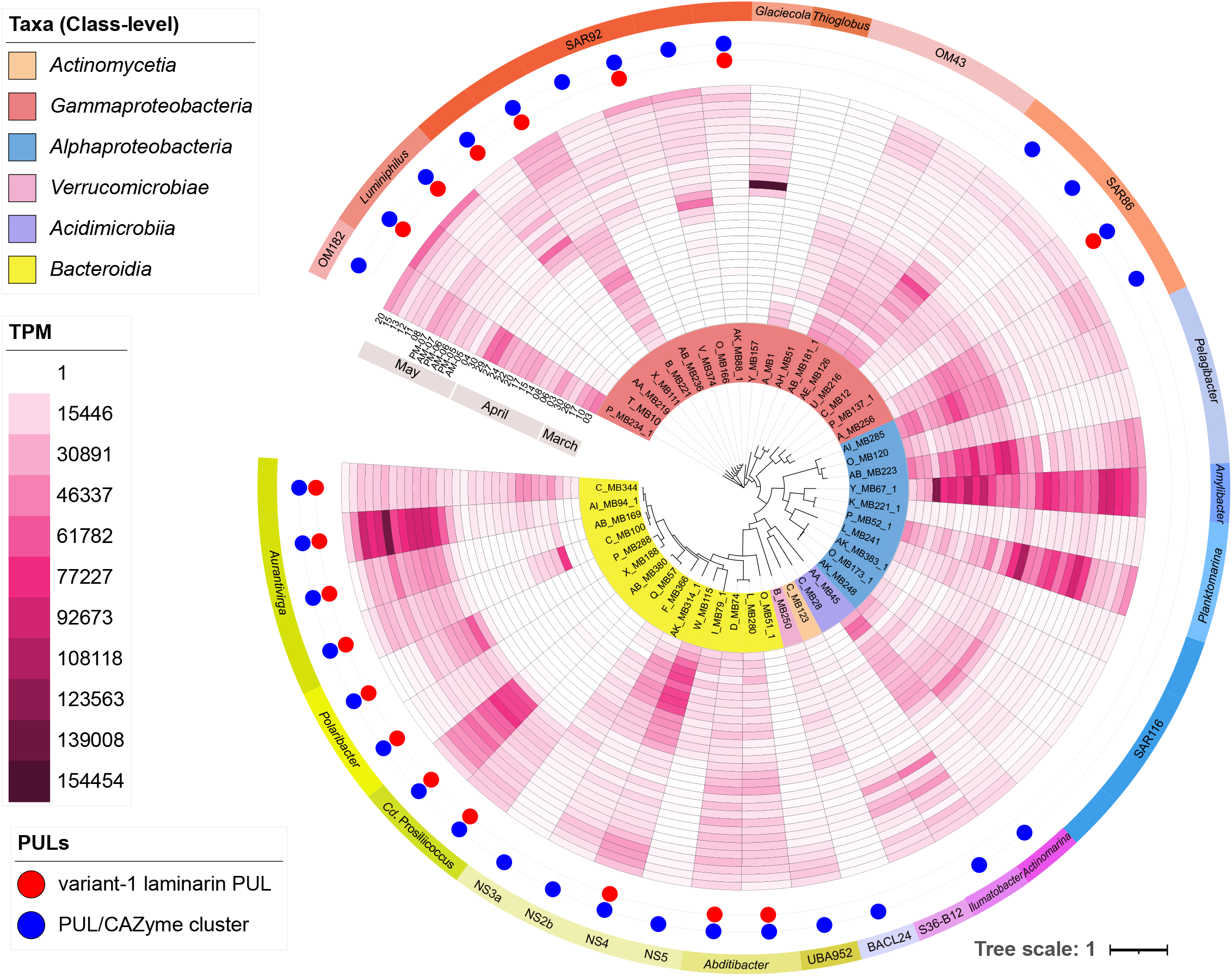
Phylogenomic tree of the top 47 expressed bacterial MAGs (created with *anvi-gen-phylogenomic-tree* of anvi’o v7) and their denominations along with their expression profiles. The expression is represented as heatmap of transcripts per million (TPM) along all sampling points from March to May 2020 (inside to outside). Genus-level clade affiliations are plotted at the outside. The outermost dots indicate presence of predicted PULs (blue) and variant 1 laminarin PULs (red), respectively. Tree annotations were done with iTOL v6.

#### - Bacteroidota

Ninety-one of the 251 representative MAGs affiliated with *Bacteroidota*, with 15 ranging among the 50 topmost expressed MAGs*. Aurantivirga* was the most active clade, with four MAGs accounting for up to 24.1% (241,055) of the TTPM. The most active of these MAGs (AI_MB94_1) exhibited highest expression during the second bloom phase after a steep rise from 1.0% (9,958) of the TTPM on April 29^th^ to 13.7% (136,921) of the TTPM on May 8^th^. The second-most active *Aurantivirga* MAG (C_MB100) exhibited maximum expression of 7.3% (72,698) of the TTMP on March 26^th^ during the *D. brightwellii* peak, indicating rapid response to proliferation of this diatom. Correlation with CARD-FISH data was high (R=0.83; p=1.03e^-007^), confirming *Aurantivirga* as prominent bloom-associated clade (**Fig. 2A**).

The second-most expressed *Bacteroidota* clade was represented by two *Cd*.Prosiliicoccus MAGs. Their combined maximum expression accounted for up to 9.5% (95,261) of the TTPM, with one MAG (AB_MB380) alone accounting for 7.5% (74,669). Expression of both MAGs was negligible during the pre- and first-bloom phases, but started to increase after April 29^th^ and for AB_MB380 peaked on May 6^th^ during the second bloom phase (**Fig. 3**). A similar pattern was observed for one of two notably expressed *Polaribacter* MAGs (P_MB288), with a peak on May 4^th^ with 6.0% (60,030) of the TTPM.

Further lower expressed *Bacteroidota* MAGs included *two Cd*. Abditibacter MAGs and four MAGs from different NS clades. The *Cd*. Abditibacter MAGs exhibited almost uniform low expression throughout March to May. In contrast, MAGs affiliating with NS5 and NS3a marine groups exhibited higher expression during the first-bloom phase, while MAGs affiliating with the NS4 and NS2b marine groups had higher expression before April 27^th^ (first phase) and after May 12^th^ (terminal phase) (**Fig. 3**).

#### - Gammaproteobacteria

Eighteen out of 78 representative gammaproteobacterial MAGs were part of the 50 expressed MAGs. A single *Glaciecola* MAG (Y_MB157) exhibited a pronounced peak on May 4^th^, making it the topmost expressed gammaproteobacterial clade with 15.4% (154,454) of the TTPM. However, its peak expression lasted for only about a day, dropped to 2.2% afterwards, and subsequently activity of *Glaciecola* Y_MB157 almost vanished with a mere 30 TPM left on May 20^th^.

Six MAGs affiliating with SAR92 exhibited high expression in the second bloom phase except MAG B_MB221, which was notably expressed throughout the entire bloom. The five other SAR92 MAGs combined contributed up to 9.3% (93,093) of the TTPM, suggesting a relevance akin to that of abundant *Bacteroidota* clades. Two *Luminiphilus* (OM60/NOR5 clade) MAGs showed only little expression during the second bloom phase, with a notable increase on the last sampling date of the terminal phase (**Fig. 3**).

In contrast, SAR86 clade MAGs showed high expression mostly during the first bloom phase, during which all four respective MAGs combined accounted for 4.0% (39,694) to 11.5% (115,285) of the TTPM. Expression decreased considerably after April 29^th^, reaching a minimum of only 0.8% (7,622) of the TTPM on May 4^th^. Similar to SAR92, expression of SAR86 increased again after May 8^th^. Also, MAGs affiliating with the OM43 clade exhibited a similar overall activity pattern.

A single MAG (A_MB1) affiliating with *Thioglobus* A (SUP05 clade) exhibited highest expression before the first bloom peak on March 26^th^, suggesting that this thiotrophic clade did not benefit from the algal bloom in the same way as other *Gammaproteobacteria*. Finally, a single MAG (P_MB234_1) affiliating with OM182 was present among the top 50 expressed MAGs, and was expressed throughout all bloom phases to varying degrees.

#### - Alphaproteobacteria

The 50 topmost expressed MAGs included ten alphaproteobacterial MAGs, with highest expression of three *Pelagibacter* (SAR11 clade) MAGs (AI_MB285, O_MB120, AB_MB223). Their collective expression ranged from 4.2% (42,104) to 16.4% (164,238) of the TTPM during the pre- and first bloom phases. Lower expression was observed for the related SAR116 clade, whose expression peaked during the first bloom phase (**Fig. 3**). An *Amylibacter* MAG (Y_MB67_1) showed high expression throughout the bloom with expression ranging from 1.7% (16,832) to 12.9% (128,976) of the TTPM. In contrast, the more active of two *Planktomarina* MAGs (P_MB52_1) was particularly active during the second bloom phase, with maximum expression of 10.9% (108,806) of the TTMP on April 29^th^.

#### - Other clades

Further highly expressed clades affiliated with *Actinobacteriota* (genera: *Actinomarina, Ilumatobacter, Cd*. Nanopelagicales) and *Verrucomicrobiota* (BACL24 clade). The top 50 expressed MAGs also contained three archaeal MAGs that all belonged to the *Poseidoniaceae* family (see supplementary text for details).

### Dissolved polysaccharides were prime targets of abundant Bacteroidota and Gammaproteobacteria

Thirty-two of the topmost 50 expressed MAGs featured expressed PULs/CAZyme clusters (**Fig. 3**). Summated PUL expressions for each predicted substrate showed highest peak expression for PULs targeting laminarin, followed by α-glucans, alginate, α-mannose/GH92-, and xylan/xylose-containing polysaccharides as well as putative porphyran. These expressions combined correlated well with algal biovolume estimates (R: 0.79; p: 1.13e^-06^) (**Fig. 4A,B**).

**Fig. 4.**
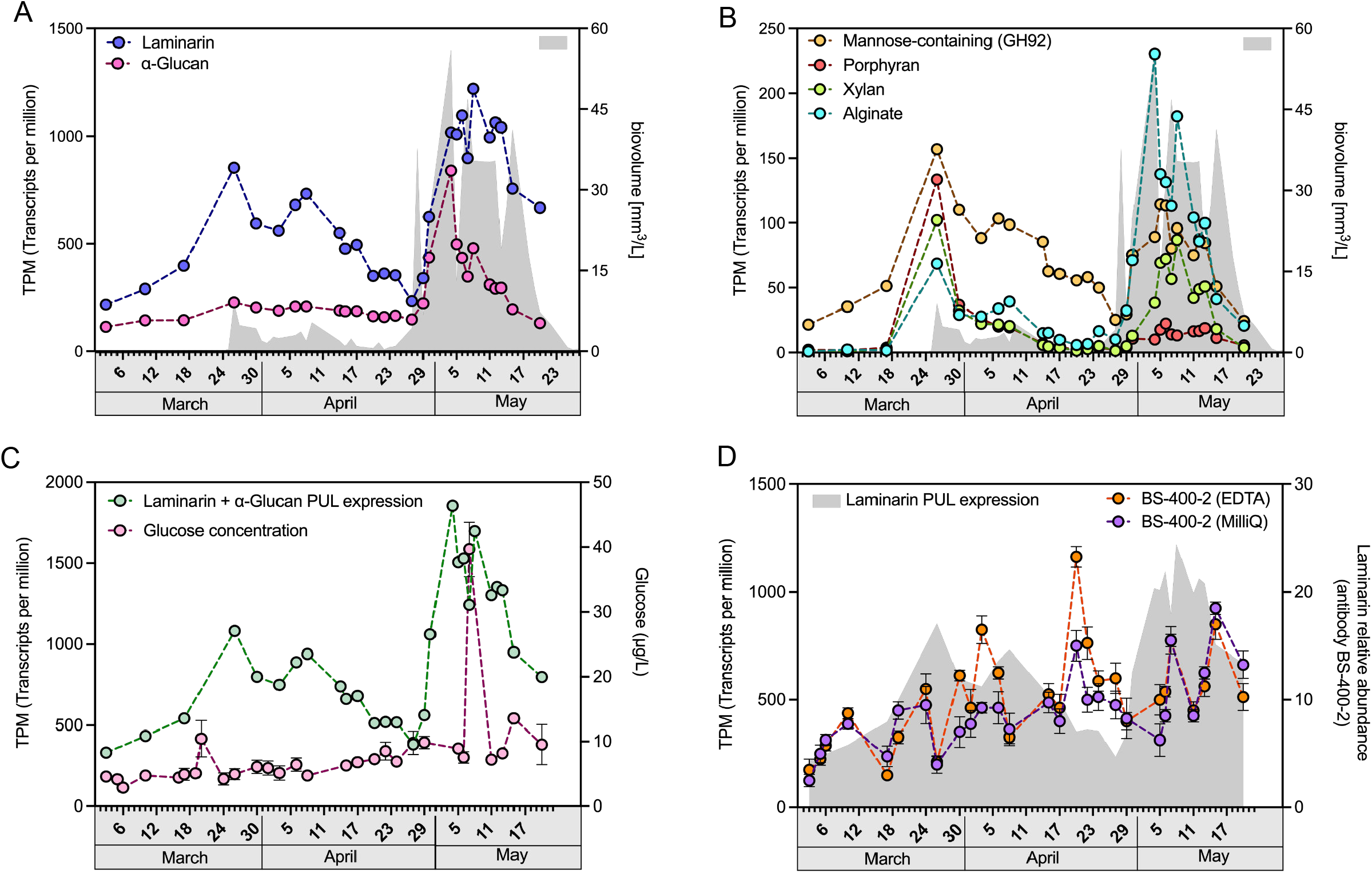
**A.** Combined expression of PULs predicted to target the two most abundant polysaccharide substrates laminarin (GH149, GH17, GH16, GH30, GH158) and α-glucans (GH13, GH65, GH31); **B.** Combined expression of PULs predicted to target less abundant polysaccharide substrates, such as mannose-rich (GH92), porphyran (GH29, GH86), xylan (GH43, GH10) and alginate (PL6, PL7, PL8, PL17). **C.** Comparison of the combined expression of PULs targeting the storage glucans laminarin and α-glucans that both consist of glucose, and measured concentrations of glucose in HMWDOM. **D.** Antibody (BS-400-2, specific for β-1,3-glucan) based measurements of dissolved laminarin extracted either with MilliQ or EDTA (lines) as compared to the sum of expressed laminarin PULs in transcripts per million (TPM; gray area).

The five bacteroidetal MAGs with the highest expression were, in descending order, *Aurantivirga* AI_MB94_1, *Cd*. Prosiliicoccus AB_MB380, *Aurantivirga* C_MB100, NS4 clade W_MB115 and *Polaribacter* P_MB288 (**Fig. 5A**). *Aurantivirga* AI_MB94_1 featured four PULs predicted to target laminarin (CAZymes: GH149-GH0-GH16_3), α-glucans (GH13), alginate (PL6-PL8-PL7) and an unspecified polysaccharide (GH3-GH130-GH18-CE2). GH0, GH16_3 and GH13 gene expression levels ranged among the top 10% in this MAG, suggesting particular active laminarin and α-glucan degradation (**Fig. 5B)**.

**Fig. 5.**
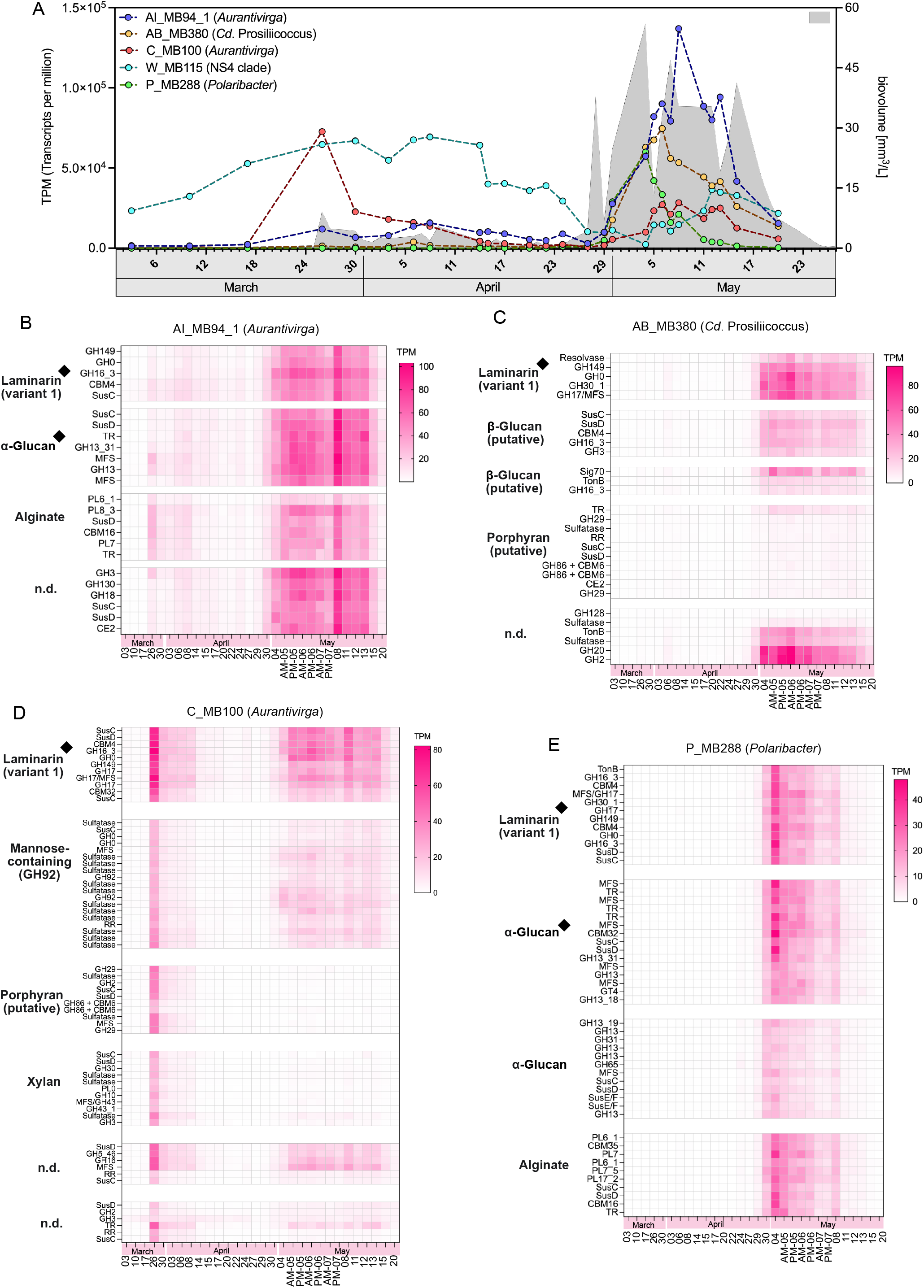
**A.** Expression of the five most expressed bacteroidetal MAGs in transcripts per million (TPM) across all sampling time-points, and total algae biovolumes (mm^3^/L) in the background (turquoise area). **B-E.** Corresponding transcriptional profiles of PULs and their predicted polysaccharide substrates: **B.** *Aurantivirga* AI_MB94_1; **C.** *Cd*. Prosiliicoccus AB_MB380; **D.** *Aurantivirga* C_MB100; **E.** *Polaribacter* P_MB288. Top expressed laminarin and α-glucan PULs are marked.

*Cd*. Prosiliicoccus MAG AB_MB380 featured three predicted expressed laminarin PULs (GH149-GH0-GH30_1-GH17, GH16_3-GH3, GH16_3). The first belonged to the previously defined variant 1 type laminarin PULs (Krüger *et al*., 2019) and exhibited the highest overall expression of these three **(Fig. 5C**). A fourth PUL, predicted to target porphyran, was not expressed. The second *Cd*. Prosiliicoccus MAG among the topmost expressed MAGs, Q_MB57, had a much weaker overall expression and featured expressed variant 1 and 2 type laminarin PULs, with slightly higher expression of the latter (**Fig. S6A**). Specialization of *Cd*. Prosiliicoccus on laminarin corroborates previous analyses (Francis *et al*., 2019).

*Polaribacter* MAG P_MB288 featured a similar set of expressed PULs as *Aurantivirga* MAG AI_MB94_1, targeting laminarin (GH149-GH0-GH17-GH16_3-GH30_1), α-glucans (GH13-GH65-GH31) and alginate (PL6-PL7-PL17). These two MAGs exhibited an almost synchronous expression peak at the onset of the second bloom phase, indicating competition for the same polysaccharide niche, in which *Aurantivirga* AI_MB94_1 notably outcompeted *Polaribacter* P_MB288 **(Fig. 5A**). Expression of the PULs in these MAGs correlated well with estimated MAG abundances and furthermore did not change relative to each other over time. Both suggest that these PULs were either unregulated or tightly co-regulated (**Fig. 5B,E**). The second *Polaribacter* MAG among the topmost expressed MAGs, X_MB188, was weaker expressed and contained expressed PULs predicted to target α-glucans and α-mannose-containing sulfated polysaccharides. Respective genes were among the topmost expressed genes in this MAG (**Fig. S6B**).

*Aurantivirga* MAGs AB_MB169 and C_MB100 were less active than AI_MB94_1, but featured more diverse expressed PULs predicted to target not only laminarin and α-glucans, but also xylan (GH43_1-GH10), a putative porphyran (GH29-GH86), a sulfated α-mannose-containing polysaccharide (GH92-sulfatases) and additional unspecified glycans (GH2-GH3; GH5-GH16) (**Figs. 5, S5**). Only the PULs targeting laminarin (variant 1) and the sulfated α-mannose-containing polysaccharide (present only in C_MB100) were notably expressed, with all glycoside hydrolases among the top 10% of expressed genes in their respective MAGs. Laminarin-targeting *Aurantivirga* C_MB100 peaked in the first bloom phase, while the *Aurantivirga* Al_MB94_1 peaked in the second bloom with highest expression of its α-glucan-targeting PUL (**Fig. 4B,D**), providing a prime example of polysaccharide niche distinction within a genus, and thereby indicating compositional changes in polysaccharide substrate availability over time.

The *Aurantivirga* MAG with the least overall expression, C_MB344, had only two expressed PULs targeting laminarin and α-glucans. In contrast to the PUL in the other three active *Aurantivirga* MAGs, these PULs did not exhibit similar expression patterns over time. While its laminarin PUL was expressed during both bloom phases, its α-glucan PUL was only expressed during the first bloom phase. This PUL featured an expressed transcriptional regulator, fortifying the view that this PUL’s expression was down regulated during the second bloom phase (**Fig. S5B**).

Additional flavobacterial MAGs affiliating with the NS4 marine group and *Cd*.Abditibacter featured variant 1 laminarin PULs (**Fig. S7, S8**). This PUL in the NS4 clade MAG W_MB115 lacked the characteristic *susCD* gene pair. NS2b MAG AK-MB314_1 lacked laminarin and α-glucan PULs, but did have PULs possibly targeting sialic acids (GH33-GH3), with GH3 among the topmost expressed genes. This MAG also featured highly expressed GH92 (α-1,2-mannosidase) genes arranged in a PUL-like structure along with *susCD* genes. In the NS3a clade MAG F_MB366, a GH13 of a putative α-glucan PUL was among the top 10% expressed genes. Finally, MAG O_MB51_1 of the *Crocinitomicaceae-*affiliating UBA952 clade featured high expression of a GH16-containing laminarin PUL during the terminal bloom phase (data not shown).

Variant 1 laminarin PULs were expressed by almost three-quarter (11/15) of the active bacteroidetal and a third (7/18) of the active gammaproteobacterial MAGs (**Fig. 3**, **Fig. 6**). Most had at least one fusion of a GH17 with an MFS transporter gene, something that was reported previously for the *Formosa* genus (Unfried *et al*., 2018). Such PULs included four of six highly expressed gammaproteobacterial SAR92 MAGs (AK_MB88, B_MB221, V_MB374, X_MB111) **(Fig. S9**), both *Luminiphilus* MAGs (AA_MB219, T_MB10) and one SAR86 (P_MB137_1) MAG, sometimes with an additional GH158 (**Figs. S9, S10**). Unlike bacteroidetal laminarin PULs, these gammaproteobacterial PULs (all SAR92) were flanked by transcriptional regulators (RNA polymerase sigma factor, ECF subfamily, *sigB*) suggesting regulation.

**Fig. 6.**
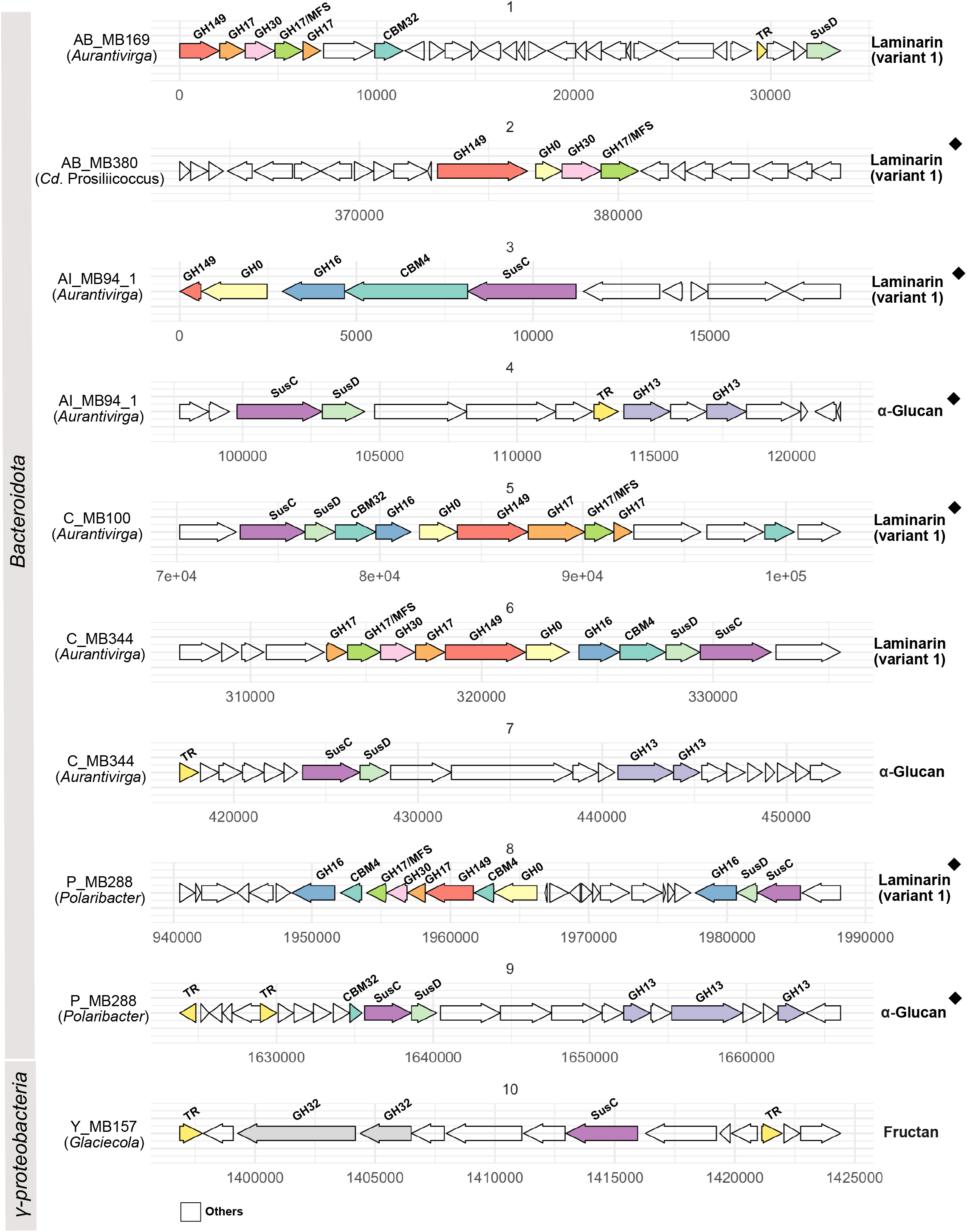
Genetic organization of highly expressed PULs predicted to target different glycans in bacteroidetal and gammaproteobacterial MAGs. PULs from top expressed MAGs mentioned in Figure 5 are marked.

In contrast to these *Gammaproteobacteria*, *Glaciecola* MAG Y_MB157 possessed seven different expressed PULs, but no variant 1 laminarin PUL. Predicted targets included α-glucans (GH13-GH31), laminarin/β-glucans (GH16_3; GH3-GH16_3; GH3), alginate (PL17_2; PL6_1) and fructans (GH32) (**Fig. S11**). Similar to many *Bacteroidota*, *Glaciecola* MAG Y_MB157 featured high expression of TonB-dependent receptors as well as expression of additional CAZymes, such as a CE4, GH13 and sugar kinase genes among the top 10% of its expressed genes.

The sole *Verrucomicrobia* MAG (B_MB250) exhibited expression of genes coding for α-L-fucosidases, arylsulfatases, and GH33 family glycoside hydrolases. Details on the predicted metabolism of *Verrucomicrobia* MAGs obtained from metagenomes off Helgoland Roads were described recently in a dedicated study (Orellana *et al*., 2021).

### Abundant storage polysaccharides played a pivotal role in the carbon flux between algal primary production, grazing and bacterial mineralization

Measurements of monosaccharide concentrations resulting from acid hydrolyzed high molecular weight DOM polysaccharides confirmed that glucose, the monomer of both laminarin and α-glucan storage polysaccharides, was the most abundant monosaccharide in the dissolved fraction during the bloom with a peak of almost 40 μg/L (**Fig. 4C)**. More importantly, changes in glucose concentrations over the course of the bloom mirrored the combined expression of predicted laminarin and α-glucan PULs over time (**Fig. 4C**). In particular, both showed a pronounced peak starting at the onset of the second bloom phase, suggesting a massive increase in the turnover of α-glucan-containing polysaccharides. Other more abundant monosaccharides comprised, in descending order, fucose (monomer in fucoidan and other fucose-containing polysaccharides), galactose (monomer in galactomannan and pectin side chains), mannose/xylose (monomers in mannans/mannose-containing and xylans/xylose-containing polysaccharides), and glucosamine (monomer in e.g. chitin) (**Table S9**). This confirmed the predicted abundant polysaccharide classes that were inferred from PUL expression data (**Fig. 4A,B**). Further monosaccharides of lesser abundance comprised rhamnose, galactosamine, glucuronic acid, arabinose and galacturonic acid (**Fig. S12**).

As expected for a diatom-dominated bloom, measurements with an anti-β-1,3-glucan antibody revealed the continuous presence of laminarin. A pronounced peak was observed on April 20^th^ (**Fig. 4D, Table S10**), coinciding with the minimum in algae biovolumes at the end of the copepod bloom (**Fig. 1B**). This increase in free laminarin most likely resulted from copepod sloppy feeding of *D. brightwellii* cells. No laminarin peak was observed at the onset of the second bloom phase, supporting the view that the strong spike in glucose at this particular time (**Fig. 4D**) resulted from available α-glucans. These α-glucans were likely bacterial in origin and resulted from increasing flagellate numbers and thus increased grazing after end of the copepod bloom (**Fig. 1B**).

## Discussion

The Helgoland 2020 spring bloom was markedly different from the blooms that we previously analyzed at this location. Top-down control by copepod grazing played a decisive role in terminating the first bloom phase, and thereby ushering in a very distinct second phase. In terms of phytoplankton composition these two phases could be considered as distinct bloom events. The bacterioplankton’s response, however, was more continuous in terms of composition and expressed gene functions and thus must be considered as response to a single biphasic substrate-producing bloom event.

During the first phase algal biovolume was dominated by the large diatom species *D. brightwellii*. Blooming *D. brightwellii* are known to inhibit proliferation of other diatom species by toxic metabolites (Rijstenbil, 1989). This might have been a contributing factor to the dominance of *D. brightwellii* during the first bloom phase. Diminishing nutrients together with strong copepod grazing likely reduced *D. brightwellii* numbers towards the end of the first bloom phase. This allowed other fast-growing diatoms, in particular large chain-forming *C. pelagica* and smaller *Chaetoceros* sp. to take over, once the copepod bloom abated. It is noteworthy in this context that blooming *C. pelagica* have been reported (i) to cope well with diminished silicate concentrations as they exist after blooms of other diatoms (Ault *et al*., 2000), and (ii) to have an adverse effect on *Calanus helgolandicus* copepod egg production (Ianora *et al*., 2008). Thus, increasing *C. pelagica* numbers at the end of the first bloom phase might have been a contributing factor to copepod bloom termination. Likewise, *Chaetoceros* sp. are able to take up silicate and other nutrients at low concentrations (Booth *et al*., 2002), and thus can thrive after blooms of other diatoms. Therefore, diminished nutrient concentrations likely played a key role in shaping algal diversity during the second bloom phase.

The unusually low algal diversity in both bloom phases allowed testing of whether blooming bacterioplankton clades responded to specific algae. Many bacterioplankton clades corresponded well with distinct diatom species. For example, during the initial *D. brightwellii* peak on 26^th^ March, bacterioplankton clades such as *Aurantivirga* (C_MB100, C_MB344), NS4, NS2b, *Amylibacter*, and SAR92 (B_MB221) showed sharp expression peaks. Likewise, *Aurantivirga* (AI_MB94_1, AB_MB169), *Polaribacter, Ulvibacter, Glaciecola*, SAR92 (except B_MB221), OM43, *Ilumatobacter* and *Cd*. Nanopelagicales activities seemed tightly coupled to the presence of *C. pelagica* and *Chaetoceros* sp. diatoms in the second bloom phase.

However, in case of the most abundant polysaccharide-degrading bacteria, the presumption of direct coupling to algae species is misleading. Laminarin PUL expression correlated well with overall diatom biomass throughout the bloom, reflecting that diatoms produce laminarin, independent of species. This was confirmed by laminarin measurements with a specific monoclonal antibody, showing that laminarin was present throughout the bloom. However, the sharp increase in glucose concentration and α-glucan PUL expression at the onset of the second phase unlikely resulted from a change in diatom species composition, but probably rather from the massive flagellate grazing at the bloom’s inflection point (**Fig. 1**). The resulting die-off of bacteria led to a copious release of bacterial storage α-glucans that allowed specialized, rivaling clades such as *Aurantivirga* AI_MB94_1 and *Polaribacter* P_MB288 to thrive. These results indicate that shifts in polysaccharide substrate availability can profoundly shape bacterioplankton community composition.

During the terminal bloom phase, expressions of all SAR86, *Pelagibacter* (AB_MB223, AI_MB285), OM43 (AB_MB181_1), SAR92 (B_MB221), NS4 (W_MB115), NS2b (AK_MB314_1) and UBA952 (O_MB51_1) MAGs increased. These clades were characterized by on overall small genome sizes (0.9-2.2 Mbp), reflecting their ability for fast growth, which enables these copiotrophs to grow quickly to acquire and utilize various low molecular weight organic algal substrates during terminal bloom phases (Ho *et al*., 2017). Moreover, relative abundances of those *Bacteroidota* that dominated the second bloom phase decreased notably during the terminal phase, indicating that they were outcompeted by fast-growing *Gammaproteobacteria* upon the massive availability of readily metabolizable algal substrates.

We have described CAZymes and PUL repertoires of phytoplankton-associated bacterioplankton clades in previous studies (e.g., Xing *et al*., 2015; Unfried *et al*., 2018; Chen *et al*., 2018; Francis *et al*., 2019; Kappelmann *et al*., 2019; Avci *et al*., 2020). Similar PUL types were also identified in the 2020 spring bloom MAGs presented here. Metatranscriptome analyses, however, revealed that the bulk of PULs that were expressed by the most active bacterioplankton clades targeted only few substrate classes, supporting hypotheses from previous genome- (Kappelmann *et al*. 2018) and metagenome-based studies (Krüger *et al*., 2019). These substrates are, from higher to lower expression, laminarin/β-glucans, α-glucans, alginates, and mannose and xylose-containing polysaccharides. In particular variant-1 laminarin PULs, as previously described in *Formosa* Hel1_33_131 (Unfried *et al*., 2018), exhibited high expression levels. Presence and expression of similar PULs in 18 abundant MAGs (**Fig. 6**) suggested a remarkable competition for laminarin among dominant bacterioplankton clades. Expression of α-glucan PULs was particularly high in all *Aurantivirga* MAG except AI_MB94_1, both *Polaribacter* MAGs, NS3a (F_MB366), and the sole *Glaciecola* MAG. During the first bloom phase, α-glucan PUL expression was highest in *Aurantivirga* MAGs AB_MB169 and C_MB344.

We observed distinct polysaccharide preferences within members of the same genus. For instance, we found high expression of GH92 α-mannosidases in *Aurantivirga* C_MB100 as well as expressed laminarin and xylan PULs, whereas *Aurantivirga* AI_MB94_1 expressed laminarin, α-glucan and alginate PULs. These preferences seemed to be either hardcoded or tightly co-regulated, because overall PUL expression patterns did not change in most *Aurantivirga* MAGs. An exception was *Aurantivirga* MAG C_MB344 that showed a notable down-regulation of its α-glucan PUL relative to its laminarin PUL during the second bloom phase (**Fig. S5B**).

Heterotrophic bacterioplankton clades which respond rapidly to phytoplankton blooms often have small genomes. For example, the top 50 expressed MAGs mostly had genome sizes between 0.8 Mbp (*Pelagibacter* and OM43 clade) and 2.9 Mbp, except *Luminiphilus* T_MB10, OM182 P_MB234_1, and *Ilumatobacter* C_MB28, whose MAGs ranged up to ~3.5 Mbp. The highest expressed MAG during the second bloom phase, *Aurantivirga* AI_MB94_1 and *Cd*. Prosiliicoccus AB_MB380 had sizes around 2.0-2.2 Mbp. Such rapid responders are often notoriously difficult to cultivate, likely because they have become auxotrophic in the process of genome streamlining.

During the last decade, we have isolated thousands of North Sea bacterial strains (Hahnke & Harder, 2013; Hahnke *et al*., 2015; Alejandre-Colomo *et al*., 2020). Despite these efforts, only two *Formosa* strains (Unfried *et al*., 2018) and one *Reinekea* strain (Hahnke *et al*., 2015; Avcı *et al*., 2017) had genomes almost identical to bacterioplankton species that were abundant during blooms. Up to now, most of the actual bloom representing bacteria remained uncultivable. Regulation is among those traits that are typically reduced during genome streamlining - *Pelagibacter ubique* being a prime example (Giovannoni *et al*., 2005). Having largely unregulated PULs without obvious transcriptional regulators could be an adaptation of *Bacteroidota* for a swift response to bloom situations and might be among the reasons, why *Bacteroidota* usually outcompete *Gammaproteobacteria* at the onset of spring phytoplankton blooms at Helgoland Roads (e.g., Teeling *et al*., 2012; Teeling *et al*., 2016).

### Study limitations and concluding remarks

A potential limitation of this study is that MAGs are more likely to reflect consensus core genomes than actual *in situ* genomes with all accessory genes. However, our previous studies have shown that in particular fast responders to phytoplankton blooms often feature an astonishing level of clonality (Avci *et al*., 2020), which is why this is unlikely to affect results. Likewise, not all polysaccharide degradation capacities are encoded in PULs, which could lead to an underestimation of bacterial polysaccharide turnover. This might affect in particular simple storage polysaccharides such as laminarin and α-glucans, which require only few genes for degradation. To make this comprehensive study feasible, we forewent sample replication in favor of a high time resolution over a long sampling period. Dense sampling over time compensates for replication to some extent and still allows for meaningful analyses (Lennon, 2011). However, it imposes limits on statistical analyses, which is why we restricted our analyses to the most pronounced major effects. Further methodological limitations, such as the specificities of the CARD-FISH probes and antibodies, are discussed in the supplementary text. In this study, we propose that organisms on higher trophic levels, such as copepods and ciliates, can influence the pool of dissolved polysaccharides available to the bacterioplankton, which we deduced from species abundances, polysaccharide concentrations, and bacterial PUL expression patterns. This hypothesis can be tested in future controlled grazing experiments.

Notwithstanding such limitations, we have a collection of powerful tools at hand for obtaining deep insights into the principles that govern bacterioplankton responses to phytoplankton blooms. Thereby we can disentangle the intricate underlying food-webs, which are key to global carbon cycling. Microbial decomposition of algal biomass is a concerted effort of both bacterial specialists and generalists with distinct substrate niches. While resource partitioning is particularly evident for polysaccharides, it also plays a role for many other abundant substrates. For example, our dataset indicated additional resource partitioning for lipids (e.g., fatty acids), proteins, compatible solutes (e.g., choline, ectoine), organosulfur compounds (e.g., DMSP), secondary metabolites (e.g., terpenoids), and C1 compounds (e.g., methanol, carbon monoxide), which are beyond the focus of this particular study. Availability and composition of algae-derived substrates therefore exert a strong influence in shaping the bacterioplankton community during phytoplankton blooms. As we show in this study, abundant storage glycans play a pivotal role in this process.

## Materials and Methods

### Sampling and filtrations

Samples were taken from March to May 2020 at the long-term ecological research site ‘Kabeltonne’ (54° 11.3’ N, 7° 54.0’ E) off the coast of Helgoland Roads (North Sea) during a diatom-dominated spring phytoplankton bloom. Seawater was sampled at 1 m depth five times a week as described previously (Teeling *et al*., 2016) for a total of 55 days.

Ten liters of seawater were filtered sequentially through 10, 3 and 0.2 μm pore-size polycarbonate filters (142 mm diameter) for bacterial biomass. The 10 and 3 μm filters retained most of the eukaryotes, larger particles and attached bacteria, while the bulk of the free-living bacteria were collected on the 0.2 μm filters. All filtrations were performed in duplicates and filters were immediately flash frozen in liquid nitrogen and kept at −80 °C until further use.

For cell counting 10 or 100 mL of unfiltered seawater were fixed with 37% formaldehyde (v/v) to a concentration of 1% (v/v) for one hour at room temperature. The fixed samples were then filtered directly onto 0.2 μm pore-sized polycarbonate filters (47 mm diameter). All filters were preserved at −80 °C until further use.

### Cell counts of total bacteria and of specific clades

Bacterial cells were stained with DAPI (4’,6-diamidino-2-phenylindole) and counted microscopically for total cell number estimates. Likewise, cells of prominent clades were stained with specific probes for CARD-FISH and counted microscopically. Both techniques were applied as described previously (Teeling *et al*., 2016). CARD-FISH probes: AUR452 (*Aurantivirga*), POL740 (*Polaribacter*), Vis6-814 (Vis6 clade including *Cd*. Abditibacter), SAR86-1245 (SAR86 clade), SAR92-627 (SAR92 clade), SAR11-mix (SAR11 clade including *Pelagibacter*) and Ros537 (Roseobacter clade). Corresponding sequences are provided in **Table S11**.

### Physicochemical and phytoplankton measurements

Physicochemical parameters such as temperature, salinity, Secchi depth, nitrate, nitrite, ammonium, phosphate, silicate as well as phytoplankton cell numbers and community composition were obtained as part of the Helgoland Roads time-series (Wiltshire *et al*., 2010; Kraberg *et al*., 2019). Phytoplankton biovolumes were determined according to Hillebrandt *et al*. 1999 (Hillebrand *et al*., 1999) in the framework of the Sylt Roads time series (see (Armonies *et al*., 2018) for details). Both, the Helogland and Sylt Roads time series are conducted by the Alfred Wegener Institute, Helmholtz Centre for Polar and Marine Research (Helgoland, Germany).

### Sequencing

Metagenome and metatranscriptome sequencing of biomass from the 0.2 μm filters was performed at the Max Planck Genome Centre (Cologne, Germany). For each time point, one filter was used for the extraction of both DNA and RNA. Isolated RNA was processed to deplete ribosomal RNAs using oligo-based customized probes (Nobori *et al*., 2018) and the remaining RNA was further used for sequencing. On the basis of chlorophyll *a* data and cell numbers, we selected 27 time points for sequencing. On three dates, samples were taken twice a day (8 AM, 9 PM), which amounts to a total of 30 metagenomes and corresponding metatranscriptomes. Sampling dates and further details are provided in **Table S12**.

### Metagenomics

#### Sequencing, assemblies and binning

Metagenome sequencing was performed on a PacBio Sequel II (Menlo Park, CA, USA) using one SMRT cell per sample in long-read HiFi sequencing mode. Details on the raw data are provided in Table S2. Assemblies were generated using Flye v2.8.3 (Kolmogorov *et al*., 2019) with options *-meta* and *-pacbio-hifi*. 16S rRNAs were extracted from metagenome assemblies using Barnnap v0.9 (Seemann, 2014) and classified using SILVAngs and the Silva database v138.1 (Pruesse *et al*., 2007; Quast *et al*., 2013). A similarity threshold of 0.97 was used to cluster the sequences for creating OTUs (operational taxonomic units).

Reconstruction of MAGs (metagenome-assembled genomes) was carried out in Anvi’o v7.0 (Eren *et al*., 2015). Initial bins were created with metabat2 (Kang *et al*.,2019) from within Anvi’o, refined by invoking the anvi-refine command, and subsequently inspected visually in terms of GC profiles and contig coverages. CheckM v1.0.18 (Parks *et al*., 2015) was used to assess MAG completeness and contamination. Since assembly and binning were performed on a per sample basis, redundant MAGs were obtained. We used dRep v3.2.0 (Olm *et al*., 2017) to dereplicate MAGs with >70% completeness and <5% contamination at 0.95 ANI (Average Nucleotide Identity). MAG phylogenomic affiliations were assigned using GTDB-Tk v1.3.0 (Chaumeil *et al*., 2019) with GTDB v202. The taxonomic affiliation of representative MAGs was further improved by extracting 16S rRNA from the MAGs and placing them in the tree using Arb v7.0 (Ludwig *et al*., 2004) and Silva v138.1.

#### Metatranscriptomics

##### - RNA read quality filtering and mapping

Metatranscriptome sequencing was performed on an Illumina HiSeq 3000 (San Diego, CA, USA). About 100 million paired-end reads (2x 150 bp) were generated per sample. Ribosomal RNA reads were filtered using SortMeRNA v3.0 (Kopylova *et al*.,2012). Remaining messenger RNA reads were quality trimmed and end repaired using the *bbduk* and *repair.sh* scripts of the BBMap v35.14 suite (https://sourceforge.net/projects/bbmap/). Reads with a minimum read length of 70 bp were subsequently mapped onto all 251 representative MAGs using Bowtie2 (Langmead & Salzberg, 2012) as a part of the SqueezeMeta v1.3.1 pipeline (Tamames & Puente-Sanchez, 2018). Mapping statistics are provided in **Table S12.**

##### - Integrated analysis

We used SqueezeMeta in merged mode for the integrated analysis of metagenomes and corresponding metatranscriptomes. Concatenated MAGs were supplied to SqueezeMeta as metagenome assembly. Contig statistics were derived using prinseq v0.20.4 (Schmieder & Edwards, 2011). RNAs were predicted using Barrnap and subsequently classified using the RDP classifier (Wang *et al*., 2007). Aragorn (Laslett & Canback, 2004) was used for the prediction of tRNA/tmRNA sequences. ORFs were predicted externally using FragGeneScan (parameters *w1* and *sanger_5*) (Rho *et al*.,2010) and searched against GenBank r239 (Clark *et al*., 2016), eggNOG v5.0 (Huerta-Cepas *et al*., 2016), KEGG r58.0 (Kanehisa & Goto, 2000), CAZy (as of 30/07/2020) (Cantarel *et al*., 2009) using Diamond (Buchfink *et al*., 2015). HMM homology searches were done using HMMER3 (Eddy, 2009) against the Pfam 33.0 database (Finn *et al*., 2016). Combined annotations were used for manual prediction of PULs and CAZyme clusters. We denominated co-localizations of CAZyme, SusCD or SusC genes as PULs, and co-localizations of CAZyme genes without *susCD* or with other transporters (e.g. of the MFS type) as CAZyme clusters. Mapping of mRNA reads against contigs (concatenated MAGs) was performed using Bowtie2 and transcripts per million (TPM) values were calculated for all MAGs of a given sample as follows: (Σ reads of sample successfully mapping to a MAG x 10^6) / (Σ lengths of contigs of the MAG x Σ number of reads in the sample). Results were visualized using the SQMtools R package.

#### Data availability

Metagenome, metatranscriptome and MAG sequence data are available from the European Nucleotide Archive (accession PRJEB52999).

#### Saccharide measurements

High molecular weight DOM was sampled using tangential flow filtration in parallel to OMICs sampling from the same water body. Samples were processed as described previously (Vidal-Melgosa *et al*., 2021) with slight modifications (see supplementary text). In brief, polysaccharides were extracted (**Table S13**) and analyzed using carbohydrate microarrays in combination with monoclonal antibodies specific for various polysaccharides. Antibody binding signal intensities were quantified with Array-Pro Analyzer 6.3 (Media Cybernetics Inc., Rockville, MD, USA).

Aliquots of the HMWDOM samples were also used for monosaccharide analysis by High-Performance Anion-Exchange Chromatography with Pulsed Amperometric Detection (HPAEC-PAD) with a DionexCarboPac PA10 column (ThermoFisher Scientific, Waltham, MA, USA) as described elsewhere (Engel & Händel, 2011; Vidal-Melgosa *et al*., 2021).

## Acknowledgements

We thank Eva Maria Brodte, Antje Wichels, Uwe Nettelmann, from the Biological Station Helgoland (BAH-AWI, Germany), the LTER team, the captains and crews from FS Aade and FS Uthörn for help with sampling, analyses, logistics, and providing lab space. We thank Marcel Huntemann (Joint Genome Institute, Berkeley, CA, USA) for assistance with MAG annotations, and Tina Trautmann (MPI Bremen, Germany) for saccharide measurements. We also thank Fengqing Wang, Mikkel Schultz Johansen (MPI Bremen) and Lilly Franzmeyer (University of Greifswald) for their help with sampling. We thank Jörg Wulf, Karl-Peter Rücknagel, Mirja Meiners, Lisa Bauer and Anja Greiser (MPI Bremen) for their technical support with CARD-FISH. The Helgoland Time Series is supported by the Helmholtz Association as an LK-II performance category program. This study was funded by the Max Planck Society and supported by the Deutsche Forschungsgemeinschaft (DFG) in the framework of the research unit FOR2406 ‘Proteogenomics of Marine Polysaccharide Utilization (POMPU)’ by grants of HT (TE 813/2-2), RA (AM 73/9-2), BF (FU 627/2-2) and TS (SCHW 595/10-2).

## Figure legends

## References

Alejandre-Colomo, C., Harder, J., Fuchs, B. M., Rosselló-Móra, R., & Amann, R. (2020). High-throughput cultivation of heterotrophic bacteria during a spring phytoplankton bloom in the North Sea. Syst Appl Microbiol, 43(2), 126066.

Alonso, C., Warnecke, F., Amann, R., & Pernthaler, J. (2007). High local and global diversity of *Flavobacteria* in marine plankton. Environ Microbiol, 9(5), 1253–1266.

Armonies, W., Asmus, H., Buschbaum, C., Lackschewitz, D., Reise, K., & Rick, J. (2018). Microscopic species make the diversity: a checklist of marine flora and fauna around the Island of Sylt in the North Sea. Helgol Mar Res, 72(1), 1–9.

Ault, T., Velzeboer, R., & Zammit, R. (2000). Influence of nutrient availability on phytoplankton growth and community structure in the Port Adelaide River, Australia:[2pt] bioassay assessment of potential nutrient limitation. Hydrobiologia, 429(1), 89–103.

Avci, B., Hahnke, R. L., Chafee, M., Fischer, T., Gruber-Vodicka, H., Tegetmeyer, H. E. et al. (2017). Genomic and physiological analyses of *‘Reinekea forsetii’* reveal a versatile opportunistic lifestyle during spring algae blooms. Environ Microbiol, 19(3), 1209–1221.

Avci, B., Krüger, K., Fuchs, B. M., Teeling, H., & Amann, R. I. (2020). Polysaccharide niche partitioning of distinct *Polaribacter* clades during North Sea spring algal blooms. ISME J, 14(6), 1369–1383.

Becker, S., Tebben, J., Coffinet, S., Wiltshire, K., Iversen, M. H., Harder, T. et al. (2020). Laminarin is a major molecule in the marine carbon cycle. Proc Natl Acad Sci U S A, 117(12), 6599–6607.

Benoiston, A. S., Ibarbalz, F. M., Bittner, L., Guidi, L., Jahn, O., Dutkiewicz, S. et al. (2017). The evolution of diatoms and their biogeochemical functions. Philos Trans R Soc Lond B Biol Sci, 372(1728), 20160397.

Booth, B.C., Larouche, P., Bélanger, S., Klein, B., Amiel, D. and Mei, Z.P. (2002). Dynamics of *Chaetoceros socialis* blooms in the North Water. Deep Sea Res, 49(22-23), 5003–5025.

Bowers, R. M., Kyrpides, N. C., Stepanauskas, R., Harmon-Smith, M., Doud, D., Reddy, T. B. K. et al. (2017). Minimum information about a single amplified genome (MISAG) and a metagenome-assembled genome (MIMAG) of bacteria and archaea. Nat Biotechnol, 35(8), 725–731.

Buchfink, B., Xie, C., & Huson, D. H. (2015). Fast and sensitive protein alignment using DIAMOND. Nat Methods, 12(1), 59–60.

Cantarel, B. L., Coutinho, P. M., Rancurel, C., Bernard, T., Lombard, V., & Henrissat, B. (2009). The Carbohydrate-Active EnZymes database (CAZy): an expert resource for Glycogenomics. Nucleic Acids Res, 37, D233–238.

Castberg, T., Larsen, A., Sandaa, R. A., Brussaard, C. P. D., Egge, J. K., Heldal, M. et al. (2001). Microbial population dynamics and diversity during a bloom of the marine coccolithophorid *Emiliania huxleyi* (Haptophyta). Mar Ecol Pro Ser, 221, 39–46.

Chaumeil, P. A., Mussig, A. J., Hugenholtz, P., & Parks, D. H. (2019). GTDB-Tk: a toolkit to classify genomes with the Genome Taxonomy Database. Bioinformatics, 36(6), 1925–1927.

Chen, J., Robb, C. S., Unfried, F., Kappelmann, L., Markert, S., Song, T. et al. (2018). Alpha-and beta-mannan utilization by marine Bacteroidetes. Environ Microbiol, 20(11), 4127–4140.

Clark, K., Karsch-Mizrachi, I., Lipman, D. J., Ostell, J., & Sayers, E. W. (2016). GenBank. Nucleic Acids Res, 44(D1), D67–72.

Eddy, S. R. (2009). A new generation of homology search tools based on probabilistic inference. Genome Inform, 23(1), 205–211.

Engel, A., & Händel, N. (2011). A novel protocol for determining the concentration and composition of sugars in particulate and in high molecular weight dissolved organic matter (HMW-DOM) in seawater. Mar Chem, 127(1-4), 180–191.

Eren, A. M., Esen, O. C., Quince, C., Vineis, J. H., Morrison, H. G., Sogin, M. L. et al. (2015). Anvi’o: an advanced analysis and visualization platform for ‘omics data. PeerJ, 3, e1319.

Falkowski, Barber, & Smetacek. (1998). Biogeochemical Controls and Feedbacks on Ocean Primary Production. Science, 281(5374), 200–207.

Field, Behrenfeld, Randerson, & Falkowski. (1998). Primary production of the biosphere: integrating terrestrial and oceanic components. Science, 281(5374), 237–240.

Finn, R. D., Coggill, P., Eberhardt, R. Y., Eddy, S. R., Mistry, J., Mitchell, A. L. et al. (2016). The Pfam protein families database: towards a more sustainable future. Nucleic Acids Res, 44(D1), D279–285.

Francis, T. B., Krüger, K., Fuchs, B. M., Teeling, H., & Amann, R. I. (2019). *Candidatus* Prosiliicoccus vernus, a spring phytoplankton bloom associated member of the *Flavobacteriaceae*. Syst Appl Microbiol, 42(1), 41–53.

Giovannoni, S. J., Tripp, H. J., Givan, S., Podar, M., Vergin, K. L., Baptista, D. et al. (2005). Genome streamlining in a cosmopolitan oceanic bacterium. Science, 309(5738), 1242–1245.

Grieb, A., Francis, T.B., Krüger, K., Orellana, L.H., Amann, R., Fuchs, B.M. (2020). *Candidatus* Abditibacter, a novel genus within the *Cryomorphaceae*, thriving in the North Sea. Syst Appl Microbiol 43(4): 126088

Hahnke, R. L., Bennke, C. M., Fuchs, B. M., Mann, A. J., Rhiel, E., Teeling, H. et al. (2015). Dilution cultivation of marine heterotrophic bacteria abundant after a spring phytoplankton bloom in the North Sea. Environ Microbiol, 17(10), 3515–3526.

Hahnke, R. L., & Harder, J. (2013). Phylogenetic diversity of *Flavobacteria* isolated from the North Sea on solid media. Syst Appl Microbiol, 36(7), 497–504.

Heins, A., Reintjes, G., Amann, R. I., & Harder, J. (2021). Particle collection in Imhoff sedimentation cones enriches both motile chemotactic and particle-attached bacteria. Front Microbiol, 12, 643730.

Hillebrand, H., Dürselen, C., Kirschtel, D., Pollingher, U., & Zohary, T. (1999). Biovolume calculation for pelagic and benthic microalgae. J Phycol, 35(2), 403–424.

Ho, A., Di Lonardo, D. P., & Bodelier, P. L. E. (2017). Revisiting life strategy concepts in environmental microbial ecology. FEMS Microbiol Ecol, 93(3), fix006.

Huerta-Cepas, J., Szklarczyk, D., Forslund, K., Cook, H., Heller, D., Walter, M. C. et al. (2016). eggNOG 4.5: a hierarchical orthology framework with improved functional annotations for eukaryotic, prokaryotic and viral sequences. Nucleic Acids Res, 44(D1), D286–293.

Ianora, A., Casotti, R., Bastianini, M., Brunet, C., d’Ippolito, G., Acri, F. et al. (2008). Low reproductive success for copepods during a bloom of the non-aldehyde-producing diatom Cerataulina pelagica in the North Adriatic Sea. Mar Ecol, 29(3), 399–410.

Kanehisa, M., & Goto, S. (2000). KEGG: kyoto encyclopedia of genes and genomes. Nucleic Acids Res, 28(1), 27–30.

Kang, D. D., Li, F., Kirton, E., Thomas, A., Egan, R., An, H. et al. (2019). MetaBAT 2: an adaptive binning algorithm for robust and efficient genome reconstruction from metagenome assemblies. PeerJ, 7, e7359.

Kappelmann, L., Krüger, K., Hehemann, J.-H., Harder, J., Markert, S., Unfried, F. et al. (2019). Polysaccharide utilization loci of North Sea *Flavobacteriia* as basis for using SusC/D-protein expression for predicting major phytoplankton glycans. ISME J, 13(1), 76–91.

Kolmogorov, M., Yuan, J., Lin, Y., & Pevzner, P. A. (2019). Assembly of long, error-prone reads using repeat graphs. Nat Biotechnol, 37(5), 540–546.

Kopylova, E., Noe, L., & Touzet, H. (2012). SortMeRNA: fast and accurate filtering of ribosomal RNAs in metatranscriptomic data. Bioinformatics, 28(24), 3211–3217.

Kraberg, A., Kieb, U., Peters, S., & Wiltshire, K. H. (2019). An updated phytoplankton check-list for the Helgoland Roads time series station with eleven new records of diatoms and dinoflagellates. Helgol Mar Res, 73(1), 1–22.

Krüger, K., Chafee, M., Francis, T. B., Del Rio, T. G., Becher, D., Schweder, T. et al. (2019). In marine *Bacteroidetes* the bulk of glycan degradation during algae blooms is mediated by few clades using a restricted set of genes. ISME J, 13(11), 2800–2816.

Langmead, B., & Salzberg, S. L. (2012). Fast gapped-read alignment with Bowtie 2. Nat Methods, 9(4), 357–359.

Laslett, D., & Canback, B. (2004). ARAGORN, a program to detect tRNA genes and tmRNA genes in nucleotide sequences. Nucleic Acids Res, 32(1), 11–16.

Lennon, J. T. (2011). Replication, lies and lesser-known truths regarding experimental design in environmental microbiology. Environ Microbiol, 13(6), 1383–1386.

Löder, M. G. J., Meunier, C., Wiltshire, K. H., Boersma, M., & Aberle, N. (2011). The role of ciliates, heterotrophic dinoflagellates and copepods in structuring spring plankton communities at Helgoland Roads, North Sea. Mar Biol, 158(7), 1551–1580.

Ludwig, W., Strunk, O., Westram, R., Richter, L., Meier, H., Yadhukumar et al. (2004). ARB: a software environment for sequence data. Nucleic Acids Res, 32(4), 1363–1371.

Mann, D. G. (1999). The species concept in diatoms. Phycologia, 38(6), 437–495.

Myklestad, S. (1974). Production of carbohydrates by marine planktonic diatoms. I. Comparison of nine different species in culture. J Exp Mar Biol Ecol, 15(3), 261–274.

Nobori, T., Velásquez, A. C., Wu, J., Kvitko, B. H., Kremer, J. M., Wang, Y. et al. (2018). Transcriptome landscape of a bacterial pathogen under plant immunity. PNAS, 115 (13), E3055–3064.

Olm, M. R., Brown, C. T., Brooks, B., & Banfield, J. F. (2017). dRep: a tool for fast and accurate genomic comparisons that enables improved genome recovery from metagenomes through de-replication. ISME J, 11(12), 2864–2868.

Orellana, L. H., Francis, T. B., Ferraro, M., Hehemann, J.-H., Fuchs, B. M., & Amann, R. I. (2021). *Verrucomicrobiota* are specialist consumers of sulfated methyl pentoses during diatom blooms. ISME J, 16, 630–641.

Parks, D. H., Imelfort, M., Skennerton, C. T., Hugenholtz, P., & Tyson, G. W. (2015). CheckM: assessing the quality of microbial genomes recovered from isolates, single cells, and metagenomes. Genome Res, 25(7), 1043–1055.

Parks, D. H., Chuvochina, M., Rinke, C., Mussig, A. J., Chaumeil, P.-A., & Hugenholtz, P. (2022). GTDB: an ongoing census of bacterial and archaeal diversity through a phylogenetically consistent, rank normalized and complete genome-based taxonomy. Nucleic Acids Res, 50(D1), D785–D794.

Pruesse, E., Quast, C., Knittel, K., Fuchs, B. M., Ludwig, W., Peplies, J. et al. (2007). SILVA: a comprehensive online resource for quality checked and aligned ribosomal RNA sequence data compatible with ARB. Nucleic Acids Res, 35(21), 7188–7196.

Quast, C., Pruesse, E., Yilmaz, P., Gerken, J., Schweer, T., Yarza, P. et al. (2013). The SILVA ribosomal RNA gene database project: improved data processing and web-based tools. Nucleic Acids Res, 41, D590–596.

Rho, M., Tang, H., & Ye, Y. (2010). FragGeneScan: predicting genes in short and error-prone reads. Nucleic Acids Res, 38(20), e191.

Rijstenbil, J. W. (1989). Competitive interaction between *Ditylum brightwellii* and *Skeletonema costatum* by toxic metabolites. Neth J Sea Res, 23(1), 23–27.

Schmieder, R., & Edwards, R. (2011). Quality control and preprocessing of metagenomic datasets. Bioinformatics, 27(6), 863–864.

Seemann, T. (2014). Prokka: rapid prokaryotic genome annotation. Bioinformatics, 30(14), 2068–2069.

Sergeev, V. N., Gerasimenko, L. M., & Zavarzin, G. A. (2002). The proterozoic history and present state of cyanobacteria. Microbiology, 71(6), 623–637.

Tamames, J., & Puente-Sanchez, F. (2018). SqueezeMeta, A Highly Portable, Fully Automatic Metagenomic Analysis Pipeline. Front Microbiol, 9, 3349.

Teeling, H., Fuchs, B. M., Becher, D., Klockow, C., Gardebrecht, A., Bennke, C. M. et al. (2012). Substrate-controlled succession of marine bacterioplankton populations induced by a phytoplankton bloom. Science, 336(6081), 608–611.

Teeling, H., Fuchs, B. M., Bennke, C. M., Krüger, K., Chafee, M., Kappelmann, L. et al. (2016). Recurring patterns in bacterioplankton dynamics during coastal spring algae blooms. eLife, 5, e11888.

Unfried, F., Becker, S., Robb, C. S., Hehemann, J. H., Markert, S., Heiden, S. E. et al. (2018). Adaptive mechanisms that provide competitive advantages to marine bacteroidetes during microalgal blooms. ISME J, 12, 2894–2906.

Vargas, C.A., Cuevas, L.A., Gonzalez, H.E., Daneri, G. (2007). Bacterial growth response to copepod grazing in aquatic ecosystems. J. Mar. Biol. Assoc. U. K. 87(3), 667–674.

Vidal-Melgosa, S., Sichert, A., Francis, T. B., Bartosik, D., Niggemann, J., Wichels, A. et al. (2021). Diatom fucan polysaccharide precipitates carbon during algal blooms. Nat Comm, 12(1), 1–13.

Wang, Q., Garrity, G. M., Tiedje, J. M., & Cole, J. R. (2007). Naive Bayesian classifier for rapid assignment of rRNA sequences into the new bacterial taxonomy. Appl Environ Microbiol, 73(16), 5261–5267.

Wiltshire, K. H., Kraberg, A., Bartsch, I., Boersma, M., Franke, H. D., Freund, J. A. et al. (2010). Helgoland Roads, North Sea: 45 Years of Change. Est Coast, 33, 295–310.

Xing, P., Hahnke, R. L., Unfried, F., Markert, S., Huang, S., Barbeyron, T. et al. (2015). Niches of two polysaccharide-degrading *Polaribacter* isolates from the North Sea during a spring diatom bloom. ISME J, 9(6), 1410–1422.

Yager, P. L., Connelly, T. L., Mortazavi, B., Wommack, K. E., Bano, N., Bauer, J. E. et al. (2001). Dynamic bacterial and viral response to an algal bloom at subzero temperatures. Limnol Oceanogr, 46(4), 790–801.

