## Supplementary Information for "Grazers affect the composition of dissolved storage glycans and thereby bacterioplankton composition during a biphasic North Sea spring algae bloom"

^*^	Corresponding authors:

**Conflict of interest**

The authors declare no conflict of interest.

**Supplementary Results**

Additional highly expressed MAGs

In the following section we describe MAGs that belonged to the 50 topmost expressed MAGs, but were not described in the main manuscript:

*- Actinobacteriota*

Three out of ten *Actinobacteriota* MAGs (AA_MB45, C_MB28, and C_MB123) ranged among the topmost 50 expressed MAGs. MAG AA_MB45 affiliated with the *Actinomarina* genus and exhibited highest expression during the first bloom phase, whereas the other two affiliating with *Ilumatobacter* (C_MB28) and *Cd*. Nanopelagicales (C_MB123) were highest expressed during the later bloom.

*- Verrucomicrobiota*

Only one out of eleven *Verrucomicrobiota* MAGs belonged to the 50 topmost expressed MAGs. This MAG (B_MB250) affiliated with the BACL24 clade (*Opitutaceae*) and showed comparatively low expression levels during the pre-bloom and first bloom phases (**Fig. 3A**).

*- Archaea*

The top 50 expressed MAGs also contained three archaeal MAGs, all of which belonged to the *Poseidoniaceae* family (class *Poseidoniia*). Expression of these MAGs was limited to the pre-bloom period and reached its maximum TPM on March 26^th^. One of these MAGs (W_MB9_1) contributed no less than 121,680 TPM (12.2%) on this date. The other two (AF_MB43, AG_MB45) also attained maximum expression on March 26^th^, contributing 38,571 (3.9%) and 25,843 (2.6%) TPM, respectively (Fig. 3B). Only minor expression was detected afterwards, indicating a negligible role of *Archaea* during the bloom (**Fig. S4**).

**Supplementary Materials and Methods**

*Sampling and processing of HMWDOM*

High molecular weight dissolved organic matter (HMWDOM) samples were obtained with a procedure described previously (Vidal-Melgosa *et al.*, 2021) with few modifications. In brief, 100 L of 0.2 μm-filtered seawater samples were concentrated to a final volume of 0.5 L using a Sartoflow Study tangential flow filtration (TFF) system (Sartorius Stedim, Göttingen, Germany). The TFF system was run with three filter cassettes with a 10 kDa cutoff (3051463901E--SW, Sartocon® Slice PESU Cassette, 0.1 m² filtration area, Sartorius Stedim). The concentrated samples containing molecules <0.2~ μm and >10 kDa were stored at -20 °C until further use. After the sampling campaign, samples were further concentrated (1 kDa cutoff), subsequently dialysed (1 kDa cutoff) and freeze dryed as described previously {Vidal-Melgosa et al., 2021, #44556}. Sampling dates: 03/03/2020, 05/03/2020, 06/03/2020, 10/03/ 20220, 17/03/2020, 19/03/2020, 24/03/2020, 26/03/2020, 30/03/2020, 01/04/2020, 03/04/2020, 06/04/2020, 08/04/2020, 15/04/2020, 17/04/2020, 20/04/2020, 22/04/2020, 24/04/2020, 27/04/2020, 29/04/2020, 05/05/2020, 06/05/2020, 07/05/2020, 11/05/2020, 13/05/2020, 15/05/2020, 20/05/2020.

*HMWDOM monosaccharide analysis by HPAEC-PAD*

For chemical glycan hydrolysis, 25 μl triplicates of each amicon-concentrated DOM sample were incubated in 1 M HCl (475 μl Milli‐Q water, 500 μl 2 M HCl) in sealed glass ampoules for 24 h at 100 °C. Afterwards, 800 μl aliquots of hydrolysate were evaporated using a RVC2-18 CD plus HCl-resistant speed‐vac (Martin Christ Gefriertrocknungsanlagen GmbH, Osterode am Harz, Germany) and resuspended in 400 μl Milli‐Q water. This corresponded to a 1:20 dilution of the Amicon-concentrated DOM samples. Monosaccharide standard solutions were prepared in an analogous fashion in 1 M HCl, evaporated and resuspended in Milli‐Q water. Monosaccharide contents were determined using High-Performance Anion-Exchange Chromatography with Pulsed Amperometric Detection (HPAEC‐PAD) with a DionexCarboPac PA10 column (Thermo Scientific) as described elsewhere (Engel & Händel, 2011).

*HMWDOM polysaccharide analysis by microarrayed antibodies*

Polysaccharide extraction of the HMWDOM samples and corresponding analysis with microarrayed polysaccharide-specific antibodies was performed as described previously (Vidal-Melgosa *et al.*, 2021). In brief, freeze-dried HMWDOM samples were homogenized, and polysaccharides were sequentially extracted with: autoclaved MilliQ water, 50 mM EDTA pH 7.5, and with 4 M NaOH containing 0.1% w/v NaBH4. Polysaccharide extracts were subsequently printed onto 0.45 µm pore-sized nitrocellulose membranes (Whatman, Little Chalfont, Buckinghamshire, UK) using a microarray robot (Sprint, Arrayjet, Roslin, UK). Due to high viscosities of the MilliQ extracts, these were first 9-fold diluted in MilliQ and then 2-fold diluted in printing buffer (55.2% glycerol, 44% water, 0.8% Triton X-100) before printing. Each sample extract was printed in four microarray spots. The arrays were probed with polysaccharide-specific antibodies conjugated to alkaline phosphatase and developed in a solution containing 5-bromo-4-chloro-3-indolylphosphate and nitro blue tetra- zolium in alkaline phosphatase buffer. Afterwards, the color signal intensity was quantified using Array-Pro Analyzer 6.3 (Media Cybernetics Inc., Rockville, MD, USA). The highest detected signal intensity, which was with the antibody JIM13 on the EDTA extract of the 6^th^ of May sample, was set to 100 and all other values were normalised accordingly. The microarray data shown correspond to probing with the mouse monoclonal antibody BS-400-2 (BioSupplies, Bundoora, Australia) and an anti-mouse secondary antibody conjugated to alkaline phosphatase (A3562, Sigma-Aldrich, St. Louis, MA, USA). Data show the mean antibody signal intensity and error bars denote the standard deviation of the four printing replicates. Further details are provided in Vidal-Melgosa *et al.* (Vidal-Melgosa *et al.*, 2021) and further information on the antibodies including initial results are summarized in **Table S13**.

**Supplementary Discussion**

*Specificities of the applied CARD-FISH probes*

CARD-FISH probes were chosen based on the abundance of microbial clades found in the metagenome dataset. It is evident, however, that the specificities of the respective probes are highly dependent on the underlying taxonomic framework. For our study we referred to the taxonomy given in the ARB-SILVA project and the respective 16S rRNA database. For the probes reported in this study we ensured that the probe specificity was matching with the taxonomic classification of the 16S rRNA genes deduced from the metagenome dataset. For some alphaproteobacterial clades like *Amylibacter* and *Planktomarina*, we failed to find suitable probes with matching specificities. This is mainly due to the high diversity within the respective clades which makes probe design almost impossible.

*Limitations of the carbohydrate microarray method*

1) The method depends on molecular probes, so we are dependent on them to detect the different glycans. When analysing a sample, we can only detect the glycans for which we have an antibody for, limiting our capacity to detect the entire glycan population of the sample (e.g. in Helgoland there may be other glycans that are present continuously during the blooms but we simply do not have antibodies that recognise their structure). We have a big library of antibodies (> 50) in our group, but it is limited.

2) The method is semi-quantitative. Although the specific concentration of the glycan epitope cannot be revealed by this method, we can detect its presence and show its relative abundance in the sample set. In this case we could detect presence of β-1,3-glucan and show its relative abundance by time. The relative abundance of the epitope is determined by the signal reported by antibody binding. The relationship between epitope concentration and antibody signal intensity has been demonstrated before, for example in (our 2021 paper - <https://doi.org/10.1038/s41467-021-21009-6> - Figure S4a) where several polysaccharides were printed including serial dilutions, which after probing resulted in antibody signals that correlated with epitope concentration.

3) Our microarray approach allows immobilisation of polysaccharides and long oligosaccharides but not of monosaccharides and short oligosaccharides (Vidal-Melgosa *et al.*, 2015; Pedersen *et al*., 2012). Therefore, we can only detect polysaccharides and long oligosaccharides which might underestimate the general polysaccharide content.

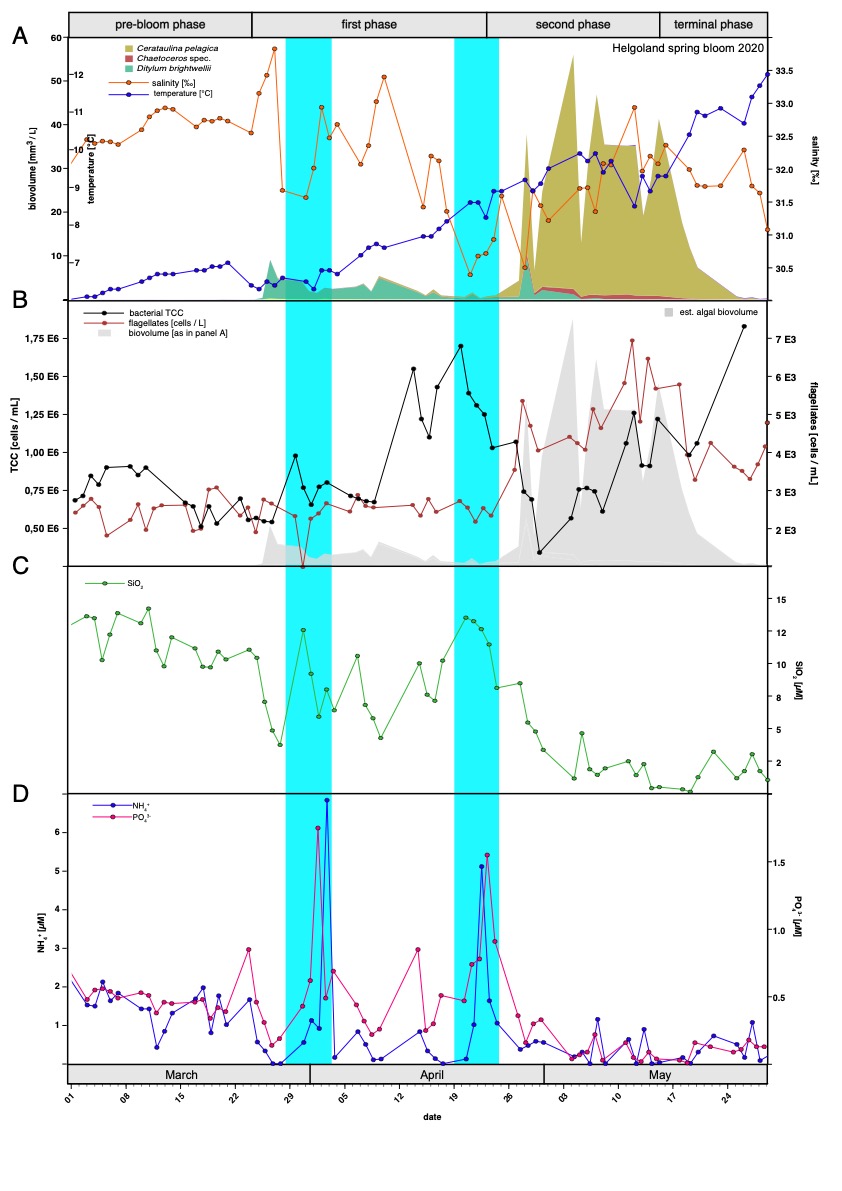

**Fig. S1** **A.** Levels of salinity and temperature measured; **B.** total bacterial cell count and flagellates; **C,** **D.** Concentration measured for inorganic nutrients such as silicate, phosphate and ammonium over a period of 2020 spring algal bloom.

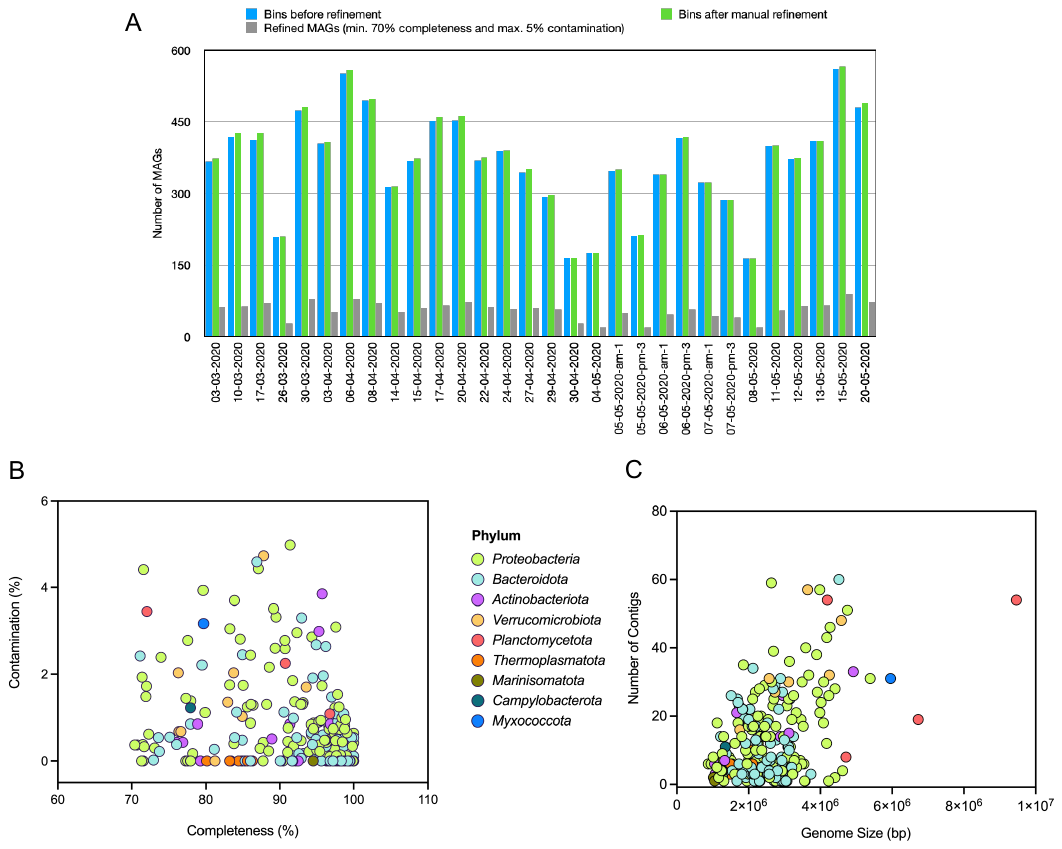

**Fig. S2** Quality of MAGs, **A.** Total number of MAGs obtained at each time-point. MAGs obtained after initial automatic binning are shown in blue color while after manual refinement using anvi-refine are shown in Green. Refined MAGs with minimum 70% completeness and maximum 5% contamination are shown in grey bars. **B.** Completeness and contamination score of refined and dereplicated 251 MAGs used in the study. **C.** Assessment of Genome size and number of contigs present in final set of dereplicated MAGs.

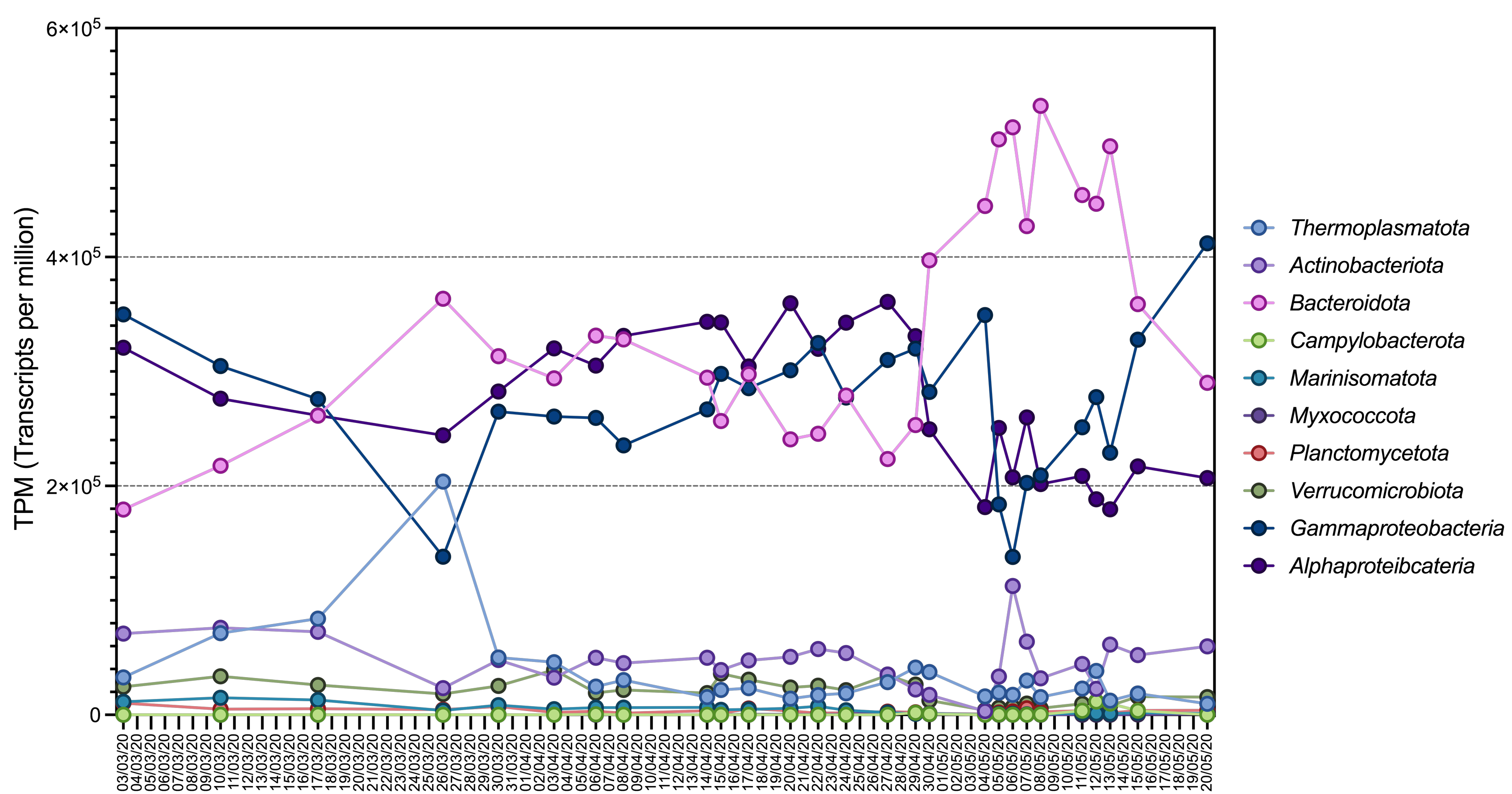

**Fig. S3** Overall expression of all clades mentioned in terms of transcripts per million during 2020 spring bloom. A clear indication of increase in TPM of *Bacteroidetes* (pink line) during the second bloom phase and decrease in TPM of *Alphaproteobacteria* (dark-blue) was observed.

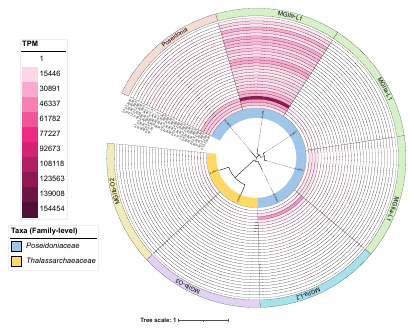

**Fig. S4** Transcription pattern of 7 Archaeal MAGs obtained in this study. All MAGs showed expression during pre-bloom phase. The MGIIa-L1 MAG (W_MB9_1) also showed transcriptional activity throughout the bloom.

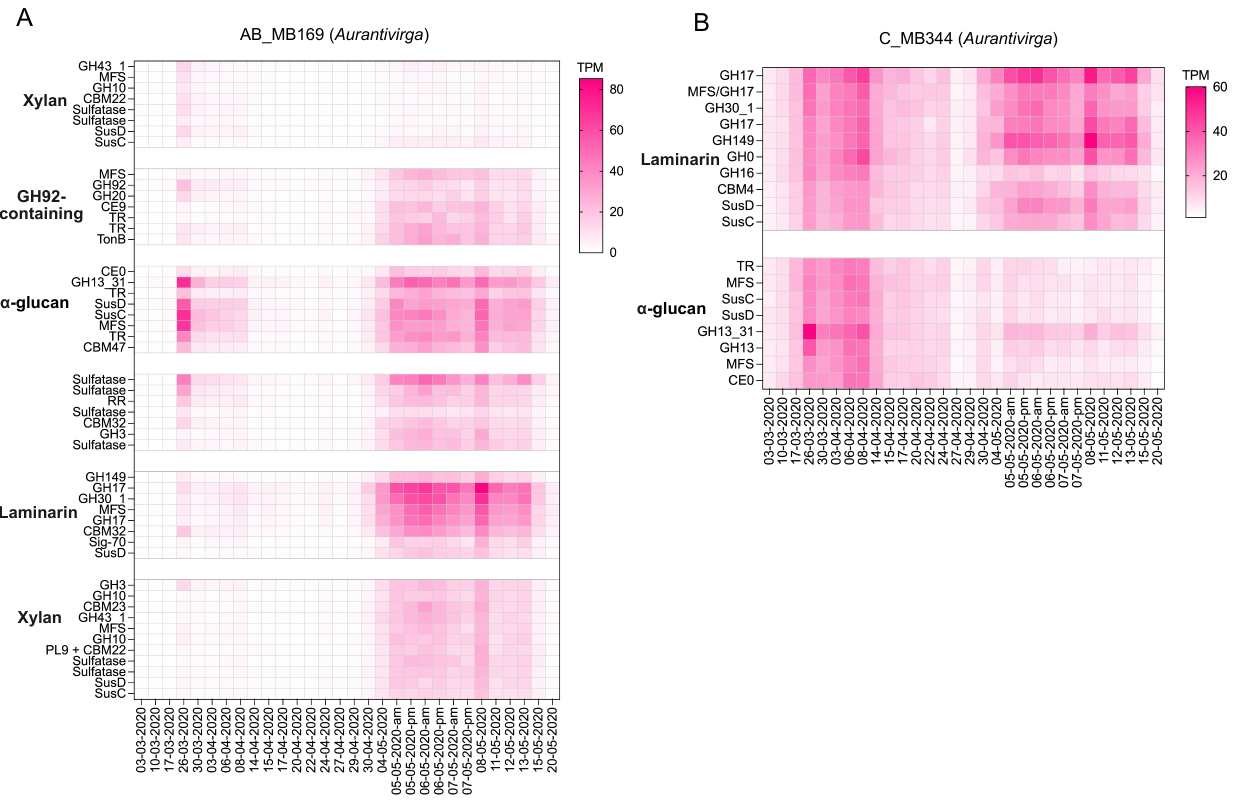

**Fig. S5** Transcriptional profiles of PULs and their predicted polysaccharide substrates in *Aurantivirga* MAG AB_MB169 and C_MB344.

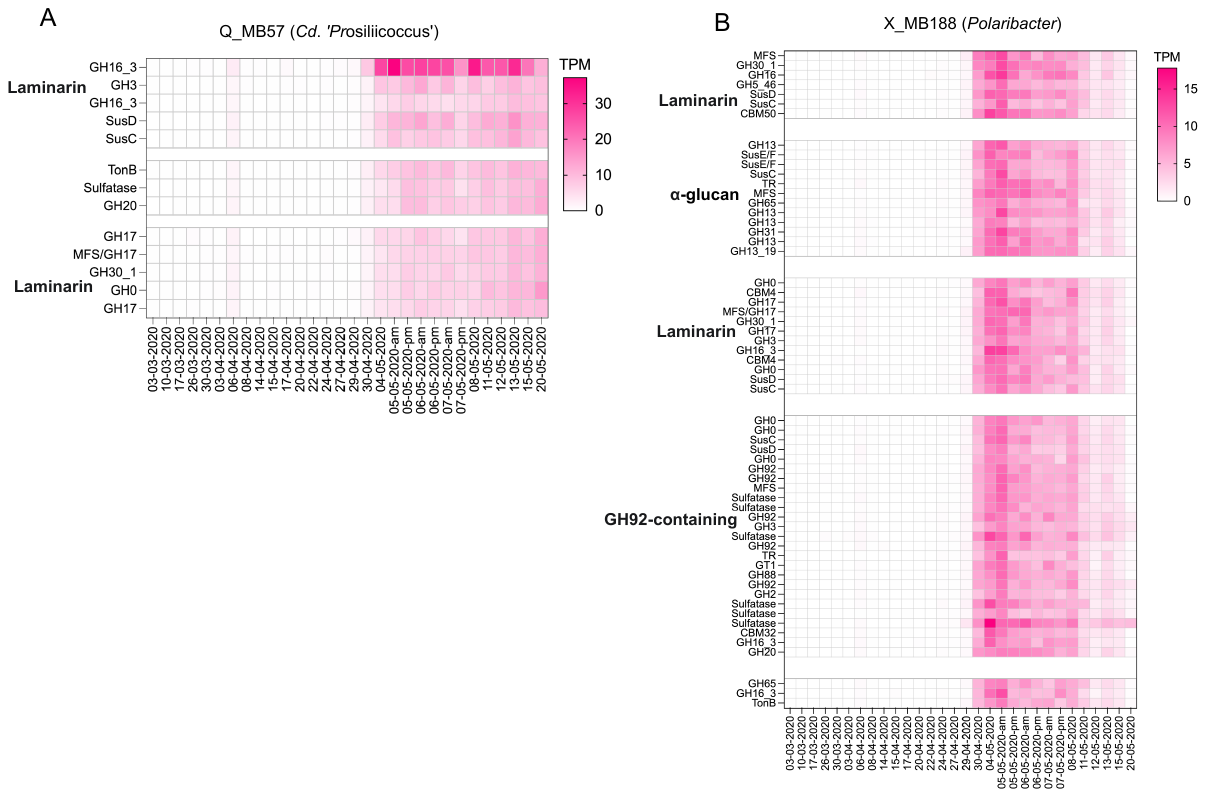

**Fig. S6** Transcriptional profiles of PULs and their predicted polysaccharide substrates in *Cd.* Prosiliicoccus MAG Q_MB57 and *Polaribacter* MAG X_MB288.

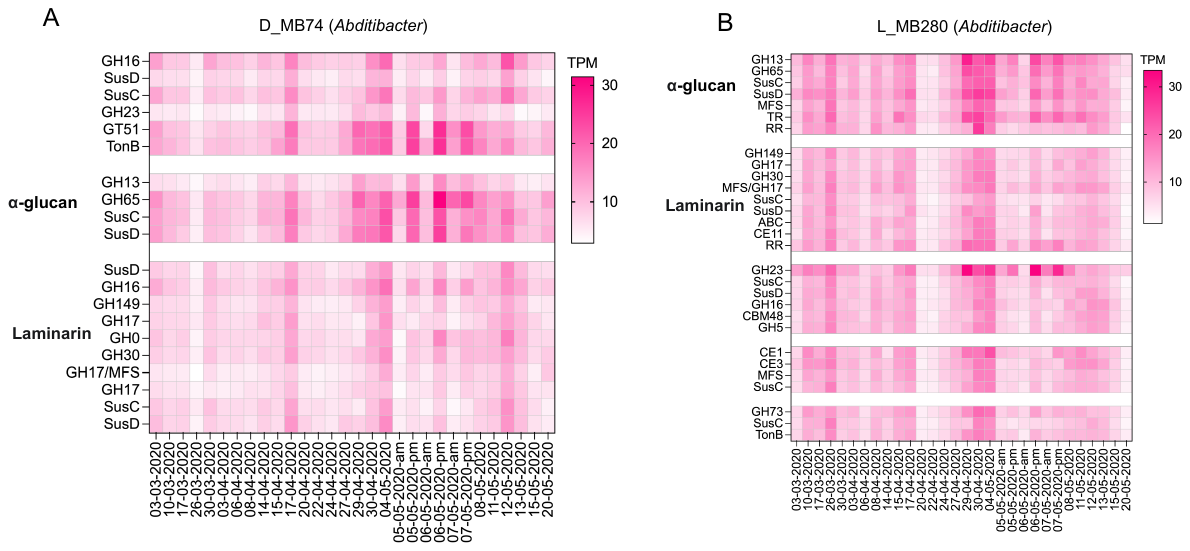

**Fig. S7** Transcriptional profiles of PULs and their predicted polysaccharide substrates in *Abditibacter* MAG D_MB74 and L_MB280.

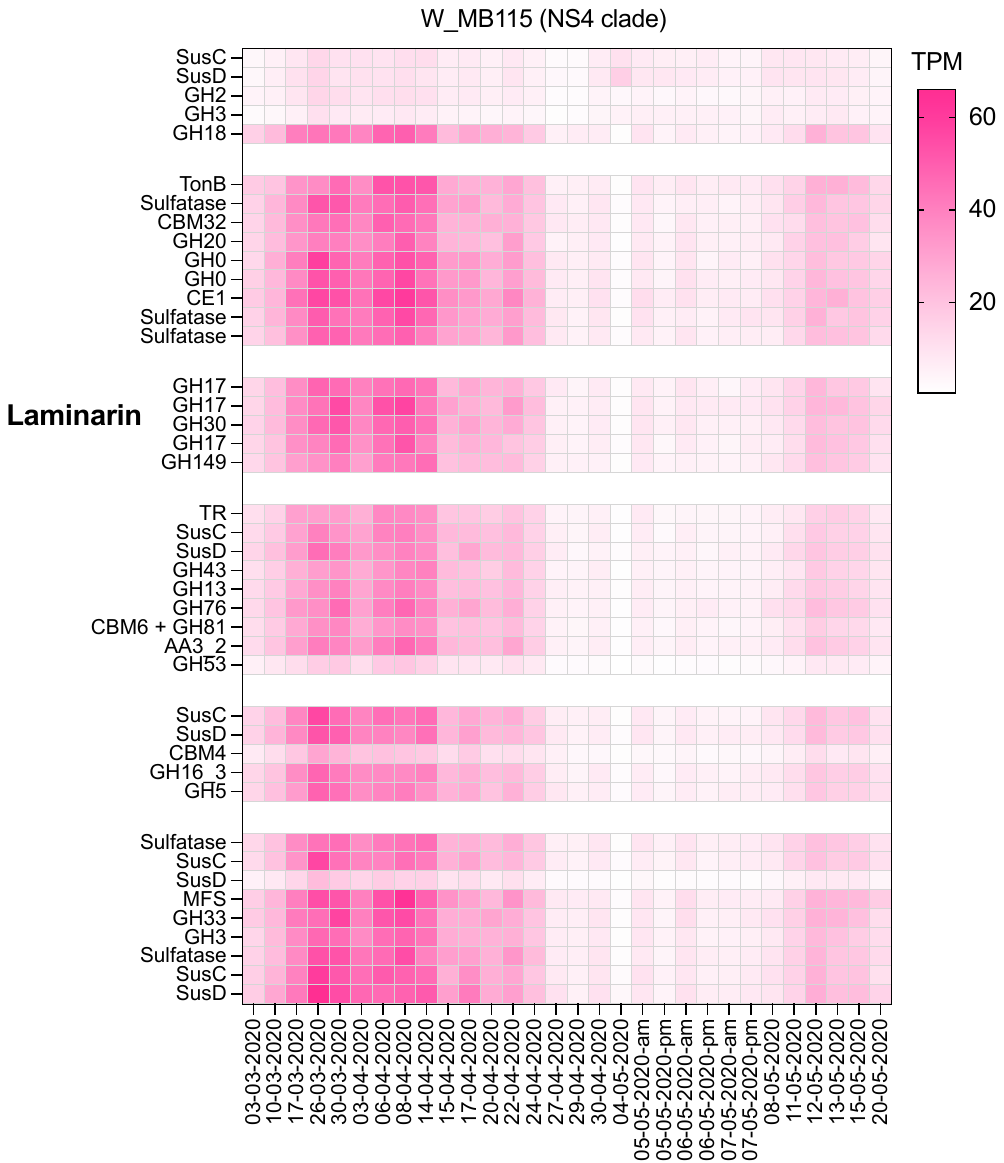

**Fig. S8** Transcriptional profiles of PULs and their predicted polysaccharide substrates in NS4 clade MAG W_MB115.

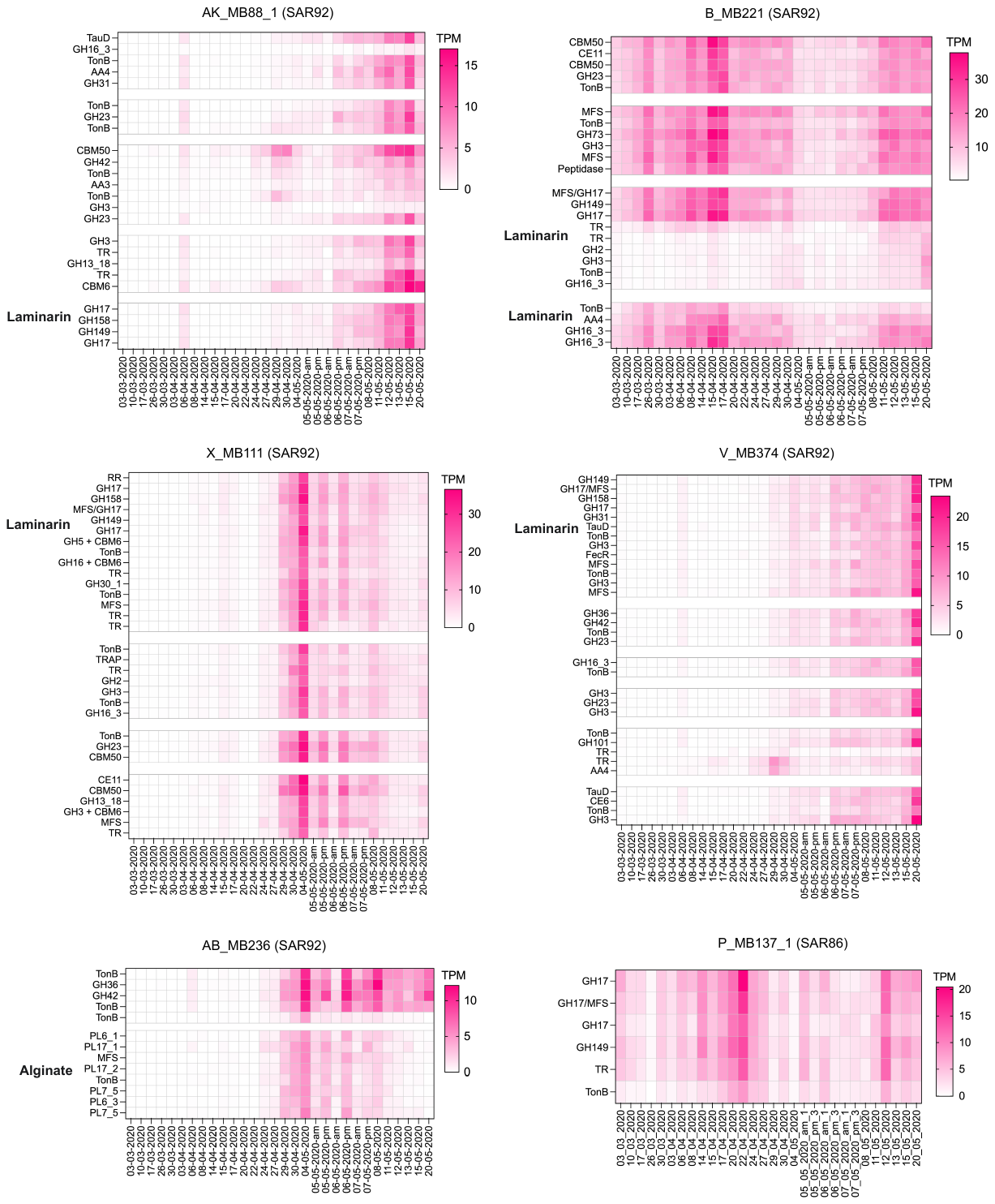

**Fig. S9** Transcriptional profiles of PULs and their predicted polysaccharide substrates in all SAR92 clade MAGs: AK_MB88_1, B_MB221, X_MB111, V_MB374, AB_MB236 and SAR86 MAG: P_MB137_1.

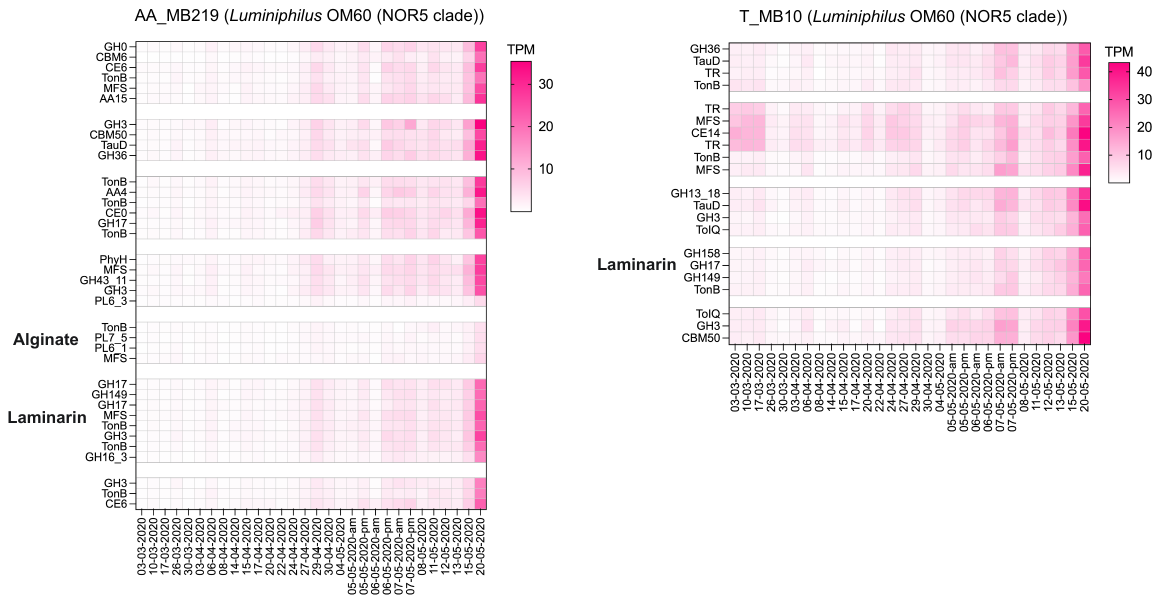

**Fig. S10** Transcriptional profiles of PULs and their predicted polysaccharide substrates in *Luminiphilus* OM60 (NOR5) clade MAGs, AA_MB219 and T_MB10.

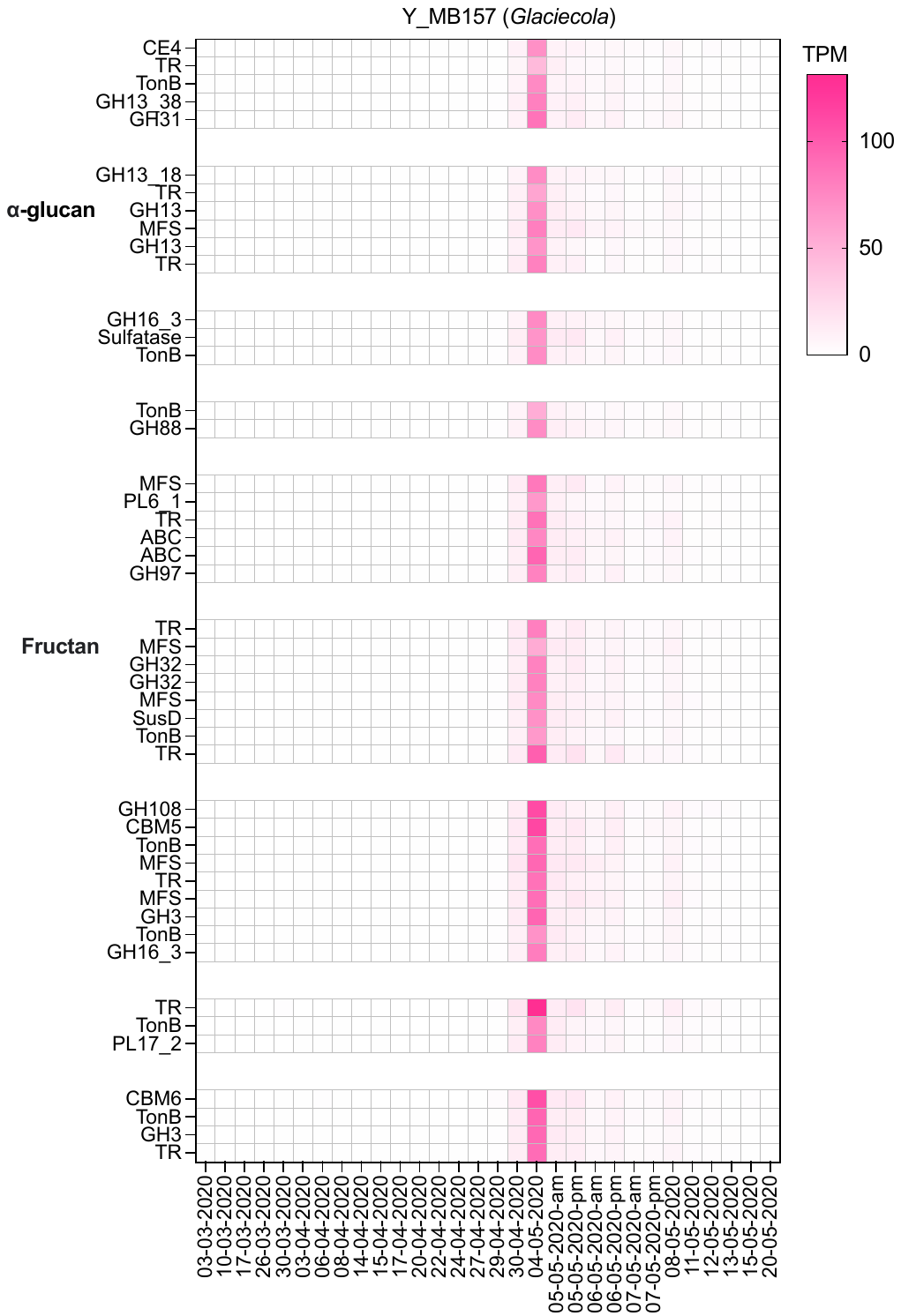

**Fig. S11** Transcriptional profiles of PULs and their predicted polysaccharide substrates in highly expressed and only Glaciecola MAG Y_MB157.

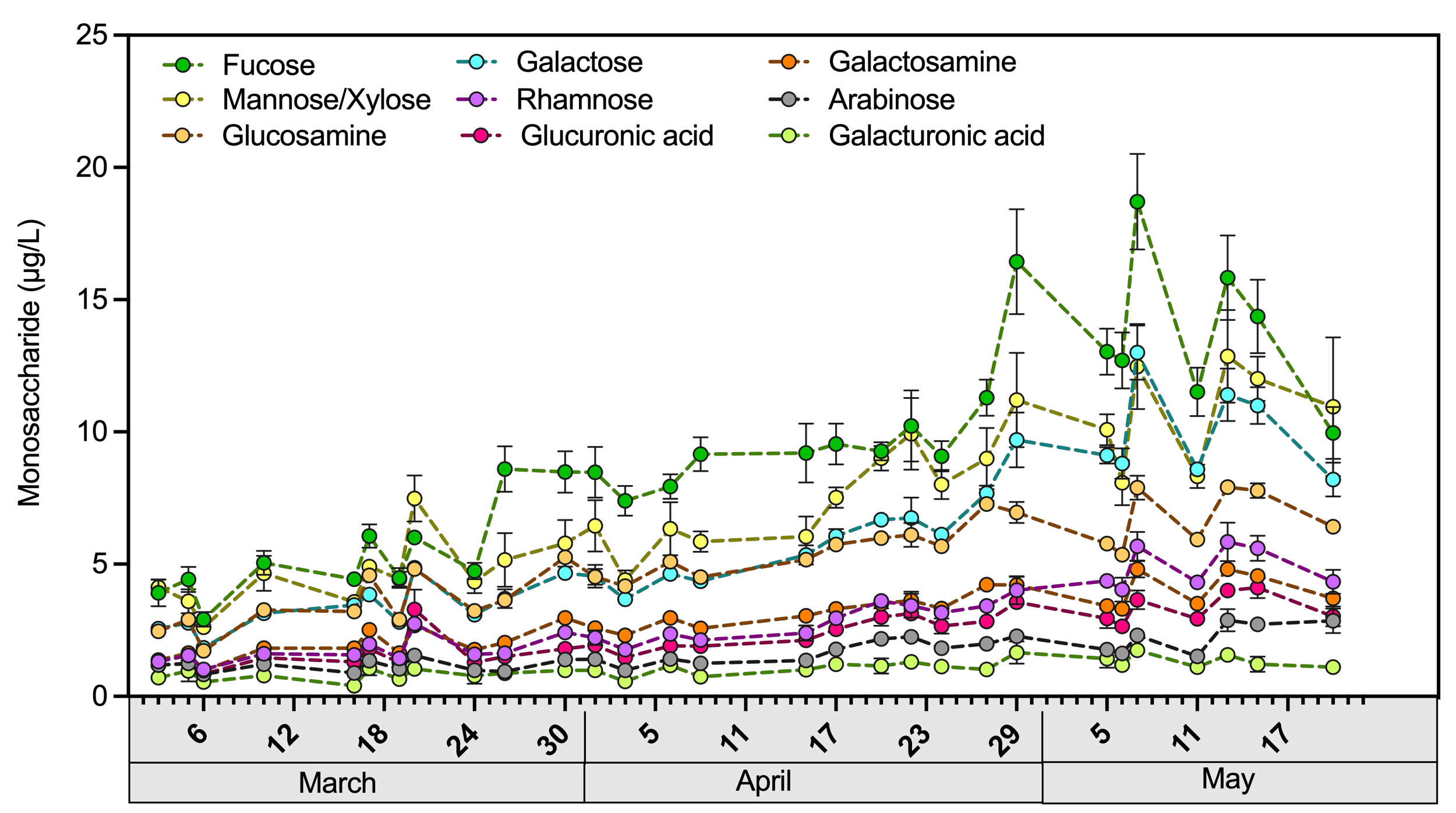

**Fig. S12** Measured concentration (in µg/L) of dissolved sugars other than glucose over all time-points.

**References**

Vidal-Melgosa, S., Sichert, A., Francis, T. B., Bartosik, D., Niggemann, J., Wichels, A. et al. (2021). Diatom fucan polysaccharide precipitates carbon during algal blooms. *Nat Comm*, *12*(1), 1-13.

Engel, A., & Händel, N. (2011). A novel protocol for determining the concentration and composition of sugars in particulate and in high molecular weight dissolved organic matter (HMW-DOM) in seawater. *Mar Chem*, *127*(1-4), 180-191.

Pedersen, H. L., Fangel, J. U., McCleary, B., Ruzanski, C., Rydahl, M. G., Ralet, M. C., Farkas, V., von Schantz, L., Marcus, S. E., Andersen, M. C., Field, R., Ohlin, M., Knox, J. P., Clausen, M. H., & Willats, W. G. (2012). Versatile high resolution oligosaccharide microarrays for plant glycobiology and cell wall research. *J Biol Chem*, *287*(47), 39429–39438.

Vidal-Melgosa, S., Pedersen, H. L., Schückel, J., Arnal, G., Dumon, C., Amby, D. B., Monrad, R. N., Westereng, B., & Willats, W. G. (2015). A new versatile microarray-based method for high throughput screening of carbohydrate-active enzymes. *J Biol Chem*, *290*(14), 9020–9036.
